# The pupal stage is a developmental window for RNA virus persistence in Drosophila

**DOI:** 10.1101/2025.11.07.687216

**Authors:** Mauro Castelló-Sanjuán, Rubén González, Alexander Bergman, Hervé Blanc, Lionel Frangeul, Jared Nigg, Maria-Carla Saleh

## Abstract

RNA viruses establish persistent infections in insects through mechanisms that are not fully understood. We focused on three positive-sense RNA viruses that naturally establish persistent infections: Drosophila A virus (DAV), Drosophila C virus (DCV), and Nora virus. We examined how these viruses interact with their hosts during development and we found that pupal metamorphosis is a critical window where virus-host immune interactions influence persistence differently across viruses. Peak viral loads and replication occur during pupation, coinciding with increased endogenous reverse transcriptase activity. Notably, reverse transcription of viral RNA genomes produces viral DNA (vDNA) forms of DAV and DCV that are first detectable during pupation and are involved in persistence. In contrast, Nora virus achieves persistence without detectable vDNA. Immune responses during pupation are virus-specific, involving suppression of RNA interference components and varied regulation of JAK-STAT signaling. After metamorphosis, DAV continues producing vDNA into adulthood while DCV shows transient vDNA production, and Nora virus bypasses vDNA production altogether. These findings point to pupation as a key developmental stage for the establishment of persistent infections through distinct viral persistence strategies.

## INTRODUCTION

RNA viruses can establish persistent infections in insects through mechanisms that remain poorly understood. These persistent infections are critical for the maintenance of viruses in nature and can have profound impacts, from the transmission of human diseases by insect vectors to economic losses in beneficial insects like bees and silkworms (Franklinos et al., 2019; Sivaprakash et al., 2025; Ullah et al., 2021). Understanding how RNA viruses achieve persistence in their hosts is therefore an important area of research for both fundamental biology and practical applications.

Persistent viral infections represent a metastable equilibrium in which viral replication is not blocked, but the physiological effects of viral infection remain below a lethal threshold (Goic & Saleh, 2012; Lee et al., 2019). Unlike acute viral infections, which are characterized by rapid viral replication that ultimately concludes in either the death of the host or the elimination of the virus, persistent infections can last for prolonged durations within the same host. Throughout this period, the virus retains the capacity for transmission to other organisms or the host’s offspring, despite the constraints imposed by the host’s immune responses and cellular mortality (Goic & Saleh, 2012).

The primary antiviral defense in insects is RNA interference (RNAi), a highly conserved pathway that provides sequence-specific protection against viral infection (Bonning & Saleh, 2021; Siomi & Siomi, 2009). In *Drosophila melanogaster*, a species in which this pathway has been extensively characterized, viral double-stranded RNA is recognized by Dicer-2 and processed into 21-nucleotide small interfering RNAs (siRNAs) (Santos et al., 2019). These siRNAs guide Argonaute-2 to cleave complementary viral RNAs, effectively suppressing viral replication (Gammon & Mello, 2015; Kim et al., 2009). This system provides rapid and specific antiviral defense, yet many RNA viruses successfully establish persistent infections despite active RNAi responses (Goic & Saleh, 2012; Randall & Griffin, 2017).

The production of viral DNA (vDNA) copies from RNA virus genomes has recently been recognized as a component of a novel antiviral mechanism in insects that facilitates viral persistence by regulating viral loads. During RNA virus infection, endogenous reverse transcriptases encoded by retrotransposons can reverse transcribe viral RNA to produce vDNA forms that often contain chimeric sequences of both viral and transposable element origin (Goic et al., 2013, 2016; Poirier et al., 2018). These vDNAs serve as long-lasting templates for antiviral siRNA production, thus amplifying and extending the RNAi response. The functional importance of this mechanism is demonstrated by the fact that preventing vDNA formation during viral infection in *D. melanogaster* or in the tiger mosquito *Aedes albopictus* results in decreased virus-specific siRNA levels and increased insect mortality (Goic et al., 2013, 2016). vDNA thus enables persistence by helping to maintain the balance between viral replication and host immune mechanisms.

vDNA formation has now been documented across diverse arthropod systems. These include Sindbis virus in *D. melanogaster* (Tassetto et al., 2017), Crimean-Congo hemorrhagic fever virus and Hazara virus in *Hyalomma*-derived tick cell lines (Salvati et al., 2021), Bombyx mori cytoplasmic polyhedrosis virus in the silkworm moth (*Bombyx mori*; Zhang et al., 2022), West Nile virus and La Crosse virus in *Aedes* cells (Nag et al., 2016), as well as chikungunya virus and dengue virus in *Aedes* mosquitoes (Goic et al., 2016). This broad phylogenetic distribution across invertebrates, including viruses from multiple families, demonstrates that the vDNA pathway represents a fundamental component of invertebrate antiviral immunity.

In *D. melanogaster*, many persistently infecting viruses are horizontally transmitted by oral acquisition of viral particles from contaminated food. Because *D. melanogaster* develops entirely on food substrates, initial encounters with viruses can occur during early developmental stages, and host–virus interactions at these stages may shape the establishment and maintenance of persistent infections. In this context, critical questions remain unanswered: How are host-virus interactions regulated during development in persistent infections? When and how is vDNA initially formed? How does host immunity enable persistence? Insect development, particularly metamorphosis, involves dramatic physiological transformations including tissue remodeling, immune system activation, and shifts in gene expression patterns (Johnston et al., 2019; Nunes et al., 2021; Zeng et al., 2025). The pupal stage represents an especially critical period during development, where larval tissues are histolyzed and adult structures are formed (Pino-Jiménez et al., 2023; Tettamanti & Casartelli, 2019; Zeng et al., 2025). In *Drosophila*, immune programming and transposable element mobilization during the pupal stage have been shown to determine adult antiviral ability (Wang et al., 2022). This developmental window could profoundly influence host-virus interactions and determine the outcome of infection, yet the interplay between viral persistence mechanisms and host development remains unexplored.

Here, we examine how three phylogenetically distinct positive-sense RNA viruses establish persistence throughout *D. melanogaster* development. Using populations persistently infected with Drosophila A virus (DAV, *Permutotetraviridae*), Drosophila C virus (DCV, *Dicistroviridae*), or Nora virus (*Picornaviridae*), we monitored viral replication dynamics, reverse transcriptase (RT) activity, and vDNA formation from embryo to adult stages. Additionally, we evaluated expression of immune genes during pupation. Our results show that pupal metamorphosis is a critical moment during viral infection, where viral persistence strategies diverge, offering new insights into the developmental regulation of antiviral immunity and the establishment of persistent infections.

## RESULTS

### Viral load and replication dynamics reveal developmental stage-specific patterns

To understand how RNA viruses establish persistence throughout *D. melanogaster* development, we analyzed viral load (presence of the virus) and replication (successful infection) dynamics over the different developmental stages in persistently mono-infected populations harboring DAV, DCV, or Nora virus infections. These infections are described to be transmitted through the oral-fecal route in the infected populations, and were previously established and confirmed to be persistent infections with individual viruses (Nigg et al., 2022, 2024). All these viruses are single-stranded positive-sense RNA viruses, with genome lengths varying between 4-12 kb (Figure 1A). To measure viral load and replication during persistent infections with each virus, adult flies persistently infected with DAV, DCV, or Nora virus were placed in clean fly food tubes for 16 h to lay eggs. For this purpose, we used 20 female flies and 20 male flies collected from standard rearing tubes used to maintain the different persistently infected lines. After the 16 h egg laying period, the F0 parental generation was discarded, and their progeny were left to develop until the adult stage. Once in the adult form, F1 adults were passed to new food every two days. Individual F1 samples from 11 developmental stages were collected and homogenized in PBS: embryo, three larval instars (L1, L2, L3), three pupal timepoints (0h, 24h, 48h post-pupation), and four adult timepoints (1, 5, 10, 20 days post-eclosion; Figure 1B). RNA extracted from a portion of the homogenate was subsequently used to measure viral load and replication levels by RT-qPCR and negative-strand specific RT-qPCR, respectively. qPCR results were analyzed using the 2^ΔCt^ method, where viral Ct values are normalized over the Ct value of a housekeeping gene (*Rp49*) for each sample.

**Figure 1.**
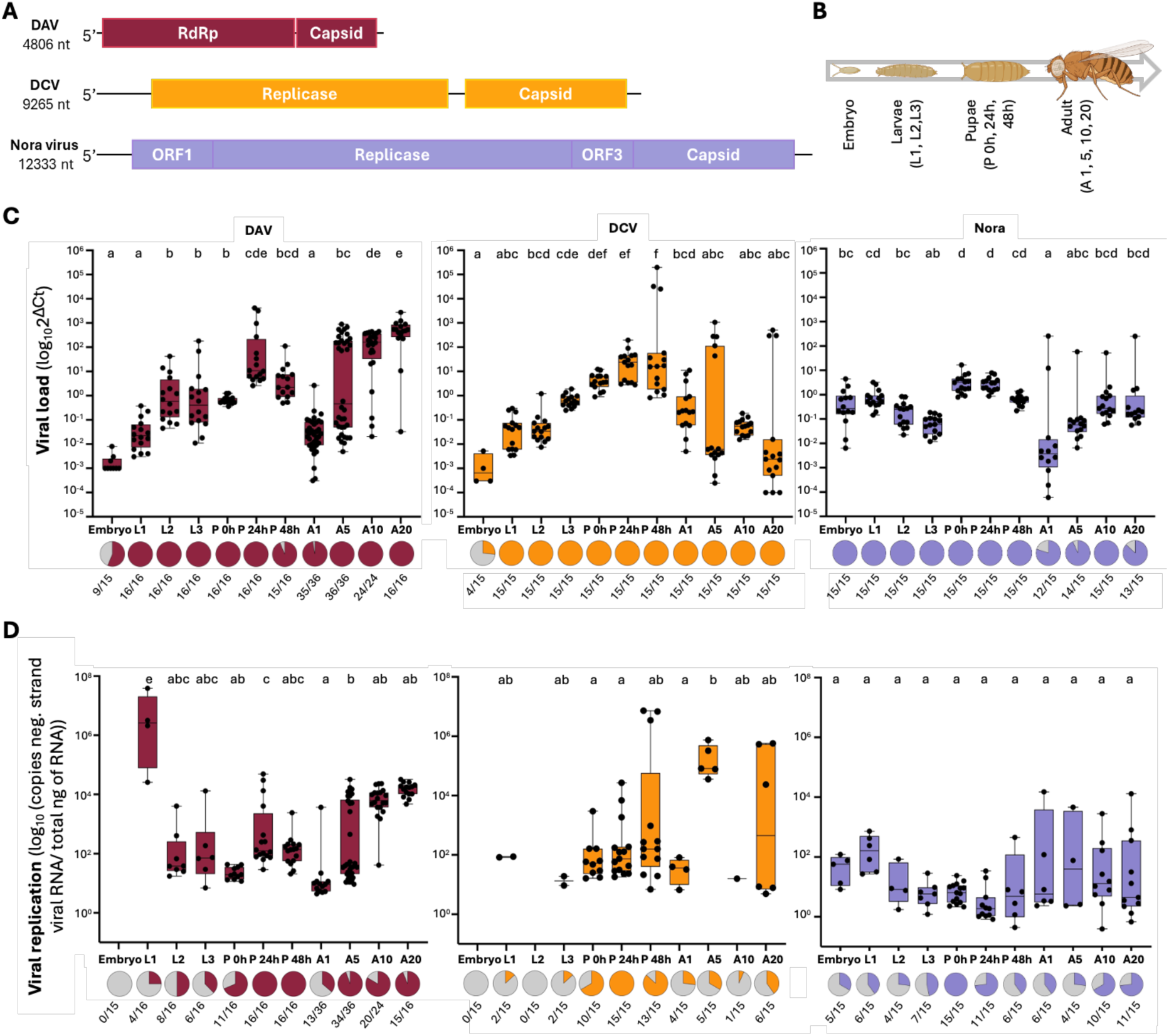
Viral load and replication in persistently mono-infected stocks. In all panels, colors indicate infection status: dark red (DAV), yellow (DCV), and light purple (Nora virus). A) Schematic representation of the DAV, DCV, and Nora virus genomes. B) Developmental stages of D. melanogaster at which samples were collected: embryo, three larval instars (L1, L2, L3), pupa at different times post-pupation (0h, 24h, 48h), and adult at various days post-eclosion (A1, A5, A10, A20). C) Viral load of DAV, DCV, and Nora virus throughout development. Y-axis depicts viral load measured as viral RNA relative to the housekeeping gene Rp49 (log_10_2^ΔCt^). X-axis indicates developmental stage of the samples. Below each developmental stage, the number of samples in which viral RNA was detected via RT-qPCR is shown. D) Viral replication of DAV, DCV, and Nora virus throughout development. Y-axis depicts viral replication measured by RT-qPCR of the negative RNA strand, quantified using a standard curve. Viral replication is expressed as log_10_ of the copies of negative-strand viral RNA per total ng of RNA in each sample. X-axis indicates developmental stage of the samples. Below each stage, the proportions of samples with detected negative-strand viral RNA (detection by RT-qPCR) are shown. Data for panels C and D were analyzed using linear mixed-effects models with developmental stage as a fixed effect and experiment as a random effect. Tukey-adjusted pairwise comparisons indicate significant differences (P < 0.05). Stages sharing lowercase letters do not differ significantly.

Following embryo hatching, viral loads increased with distinct virus-specific patterns. DAV and DCV loads remained relatively low throughout the larval stages before sharply increasing during pupation, peaking at 24 hours post-pupation (Figure 1C) then declining toward adult emergence (Figure 1C). Nora virus displayed a different trajectory, maintaining stable loads throughout embryonic and larval development (2^ΔCt^ values: 10^-0.55 ±0.79^ for the embryos, 10^-1.2±0.38^ for L3 larvae), before peaking at pupation onset (P 0h with 2^ΔCt^ values: 10^0.44±0.37^; Figure 1C). Adult-stage dynamics revealed further virus-specific patterns. DAV established robust infections with progressively increasing loads, reaching 2^ΔCt^ values of 10^2.39±1.15^ by 20 days post-eclosion (dpe; Figure 1C). DCV accumulation exhibited a bimodal distribution. While all pupae showed high viral loads and active replication, adult populations split into high viral RNA level (10^2^) and low viral RNA level (10^-3^) subgroups, with most individuals falling into the low viral RNA level category (Figure 1C). Nora virus loads accumulated steadily to 2^ΔCt^ values of 10^-0.44±0.96^ by 20 dpe (Figure 1C).

Viral replication dynamics, measured by negative-strand specific RT-qPCR, revealed that detectable replication is generally more prevalent for all three viruses during pupal stages, despite their different infection trajectories (Figure 1D). DAV and DCV first showed detectable replication in L1 larvae, with both viruses reaching 100% detection prevalence at 24 h post-pupation. However, the infection patterns of these viruses diverged during the adult stages. DAV replication declined to 36% detection prevalence post-eclosion (13/36 adults), then rebounded to 94% at 5 dpe (34/36 adults), before decreasing again to 83% by 10 dpe (20/24 adults; 10^3.73 ± 0.61^ copies of negative strand RNA/total ng of RNA; Figure 1D). DCV replication dropped sharply from 100% detection prevalence in pupae to just 27% at 1 dpe (4/15 adults). At 20 dpe, DCV replication had 40% detection prevalence, with detectable replication levels showing a bimodal distribution ranging from 10^6^ to 10^1^ copies of negative strand RNA/total ng of RNA (Figure 1D). Consistent with vertical transmission, Nora virus showed replication from the embryonic stage onward, achieving universal detection at 0 h post-pupation (15/15 samples). Unlike DAV and DCV, Nora virus maintained stable replication in adults, with replication detected in 67% (10/15) and 73% (11/15) of samples at 10 and 20 dpe, respectively, at consistent levels (10^1.34 ± 1.11^ for 10 dpe; 10^1.28 ± 1.35^ copies of negative strand RNA/total ng of RNA for 20 dpe; Figure 1D). Viral replication dynamics are virus-dependent, with varying levels of replication over the adult timepoints. This indicates that each virus interacts with the host in a specific manner, highlighting the complexity of persistent viral infections.

To understand how infected organisms manage viral replication and viral load, we can investigate correlations between these two factors. This approach can offer valuable insights into the processes that occur within persistently infected organisms. Specifically, a positive correlation between viral load and replication may suggest high tolerance or a lack of viral load control, which could potentially result in mortality in flies. The uncoupling of viral replication from the expected accumulation of viral load would indicate mechanisms of viral load control.

When analyzing the correlation between viral load and replication, we observed a positive correlation for DAV-and DCV-infected samples during pupation (Supplementary Figure 2). DAV-infected samples show this correlation during 24 and 48 hours post-pupation, as well as at 20 dpe in adults. For DCV-infected samples, the positive correlation between viral replication and viral load occurs throughout all of pupation, until and including the day of adult emergence, and again at 20 dpe in adults. In the case of Nora virus infection, we observe no such correlation between replication and viral load throughout development (Supplementary Figure 2). This is consistent with the effects observed in Nigg et al., 2024, where mortality in DAV and DCV-infected stocks was higher than those infected with Nora virus. This suggests that the modulation of replication and viral load during Nora virus infection allows for viral persistence with reduced virulence (lower pathogenicity), rather than necessarily implying a direct fitness cost to the host.

The detection of viral RNA in the embryo samples by qPCR suggested vertical viral transmission. To test this, we performed further analysis. Embryos collected from the infected lines were surface-sterilized (50% bleach treatment for 10 min). While viral RNA was detected by RT-qPCR for all three viruses, only Nora virus showed consistent detection across all embryos tested (15/15), compared to variable and near-detection-limit levels for DAV and DCV (Figure 1C). To determine whether this represented active infection or residual RNA, we performed RT-PCR targeting multiple genomic regions. All three target regions amplified successfully from Nora virus-infected embryos, confirming the presence of complete viral genomes (Supplementary Figure 1). In contrast, no additional regions amplified from DAV or DCV samples, indicating only RNA fragments remained. This pattern was corroborated by detection of negative-strand RNA (indicative of active replication) exclusively in Nora virus-infected embryos (Figure 1D), demonstrating vertical transmission for Nora virus but not DAV or DCV, a transmission route not previously documented for this virus.

### Viral infection modulates reverse transcriptase activity during development

For all three viruses, we observed that infections begin in early development and persist into adulthood. Since vDNA produced by endogenous RT activity is linked to persistent viral infection in adult arthropods (Goic et al., 2016; Poirier et al., 2018; Tassetto et al., 2017), we investigated whether RT activity was detectable during infections initiated in early host development and whether viral infection influences RT activity levels. The synthesis of vDNA from RNA virus genomes requires RT activity, yet RNA viruses do not encode this enzyme (Goic et al., 2013, 2016). To investigate whether viral infection influences endogenous RT activity levels, we first needed to confirm that RT activity is present in *D. melanogaster* throughout development. We employed an RT assay previously adapted for whole-insect conditions (Bergman et al., 2025; Pyra et al., 1994; Wu et al., 2021), where reverse transcription of exogenous MS2 phage RNA by RT activity in protein homogenates is quantified relative to 1 unit of SuperScript II as a 100% reference (further details in Materials and Methods). Additional results showed that our total protein working concentration of 1 ng/μl is well within the linear range of detection of the assay for *Drosophila* homogenates from different developmental stages (Supplementary Figure 3).

Our results demonstrate that RT activity is detectable across all developmental stages of *D. melanogaster* (Supplementary Figure 4). In uninfected flies, RT activity is highest during the embryonic stage (0.036 ± 0.026%), in first instar larvae (0.037 ± 0.051%), at 0 h post-pupation (0.050 ± 0.081%), and in 1 dpe adults (0.064 ± 0.091%).

Viral infection significantly altered these patterns. Most notably, all three viruses induced a significant increase in RT activity compared to uninfected controls at 24 hours post-pupation (DAV: 0.140 ± 0.093%; DCV: 0.257 ± 0.214%; Nora: 0.210 ± 0.141%; Figure 2, Supplementary Figure 5). Additionally, embryos from DCV-and Nora virus-infected stocks exhibited elevated RT activity (DCV: 6.767 ± 5.750%; Nora: 1.667 ± 0.816%) (Figure 2). During larval development, Nora virus infection increased RT levels across all three instars, while DCV infection elevated activity only in L3 larvae (0.021 ± 0.0009%; Figure 2). Virus-specific modulations included sustained elevation of RT activity at 48 h post-pupation for Nora virus (0.096 ± 0.068%) and suppressed RT activity at 1 dpe in DAV-infected adults (0.015 ± 0.012%; Supplementary Figure 5).

**Figure 2.**
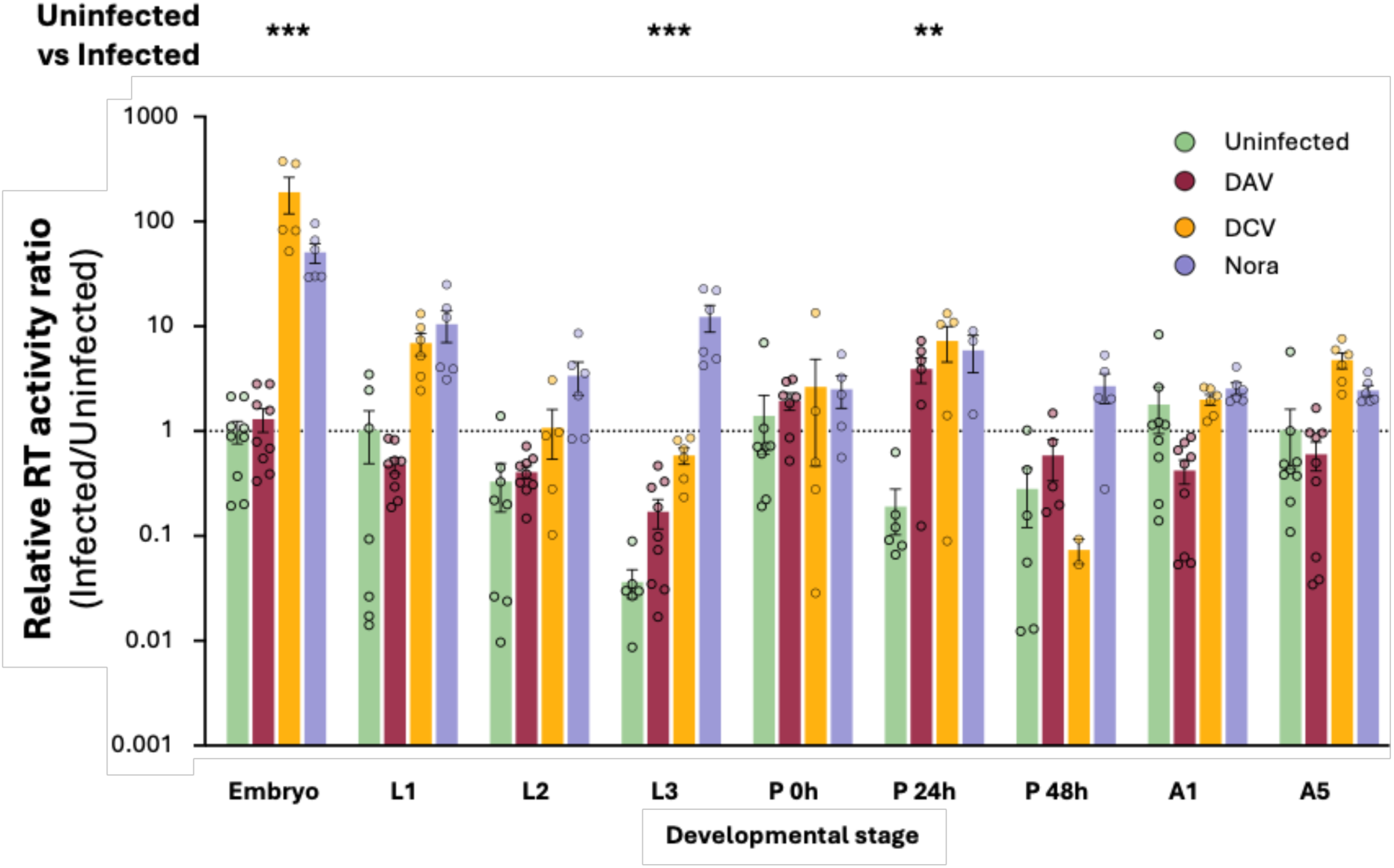
RT activity throughout development in persistently infected or uninfected flies. Y axis depicts ratio between the RT activity value of each sample and the median RT activity value of the uninfected embryos. The median RT activity value of the uninfected embryos was set to 1 and is indicated by the dotted black line. Bar colors indicate infection status: green (uninfected), dark red (DAV), yellow (DCV), and light purple (Nora virus). RT activity in uninfected flies was compared to the collective RT activity in infected flies (DAV, DCV, and Nora virus-infected) for each stage. This analysis considers two groups at each stage: uninfected and infected (DAV, DCV, and Nora virus samples). Statistical analysis was performed using a generalized linear mixed-effects model (GLMM, Gamma distribution) with infection status (Uninfected vs. Infected) and developmental stage as fixed effects, experiment and virus (DAV, DCV, Nora) as random effects. Post hoc pairwise comparisons (emmeans) tested differences between Uninfected and Infected within each developmental stage, and significance is indicated by asterisks (P < 0.05, ** P < 0.01, *** P < 0.001). Pairwise comparisons for each virus and stage-matched uninfected controls are shown in Supplementary Figure 5.

These results demonstrate that infection by DAV, DCV, or Nora virus significantly modulates RT activity across *D. melanogaster* development, with all three viruses consistently inducing RT activity during pupation, but also at earlier stages for DCV and Nora virus (Supplementary Figure 5).

### vDNA production starts during pupation and is only detected for DAV and DCV

vDNA synthesized from RNA virus genomes serve as markers of host-virus interactions and contributes to antiviral defense in insects (Goic et al., 2013; Poirier et al., 2018). To determine when vDNA is produced during development, we designed primers targeting∼200-bp amplicons spanning the entire viral genomes. Initial screening on DNA from mono-infected fly populations identified several positive amplicons for DAV and DCV. For each virus, we selected three primer pairs (designated A, B, and C) for validation by Sanger sequencing to confirm viral origin (Figure 3A). We did not detect Nora virus vDNA using this approach. For samples infected with DAV and DCV, we extracted DNA from individual fly homogenates. These were the remaining portions of the same homogenates initially used for RNA extraction to measure viral load and replication levels. We then prepared virus-and developmental stage-specific DNA pools (four individuals per pool) for detection of vDNA by PCR (Figure 3A). vDNA was first detected during pupation for both viruses and is subsequently detected again in the adult stages (Figure 3A).

**Figure 3.**
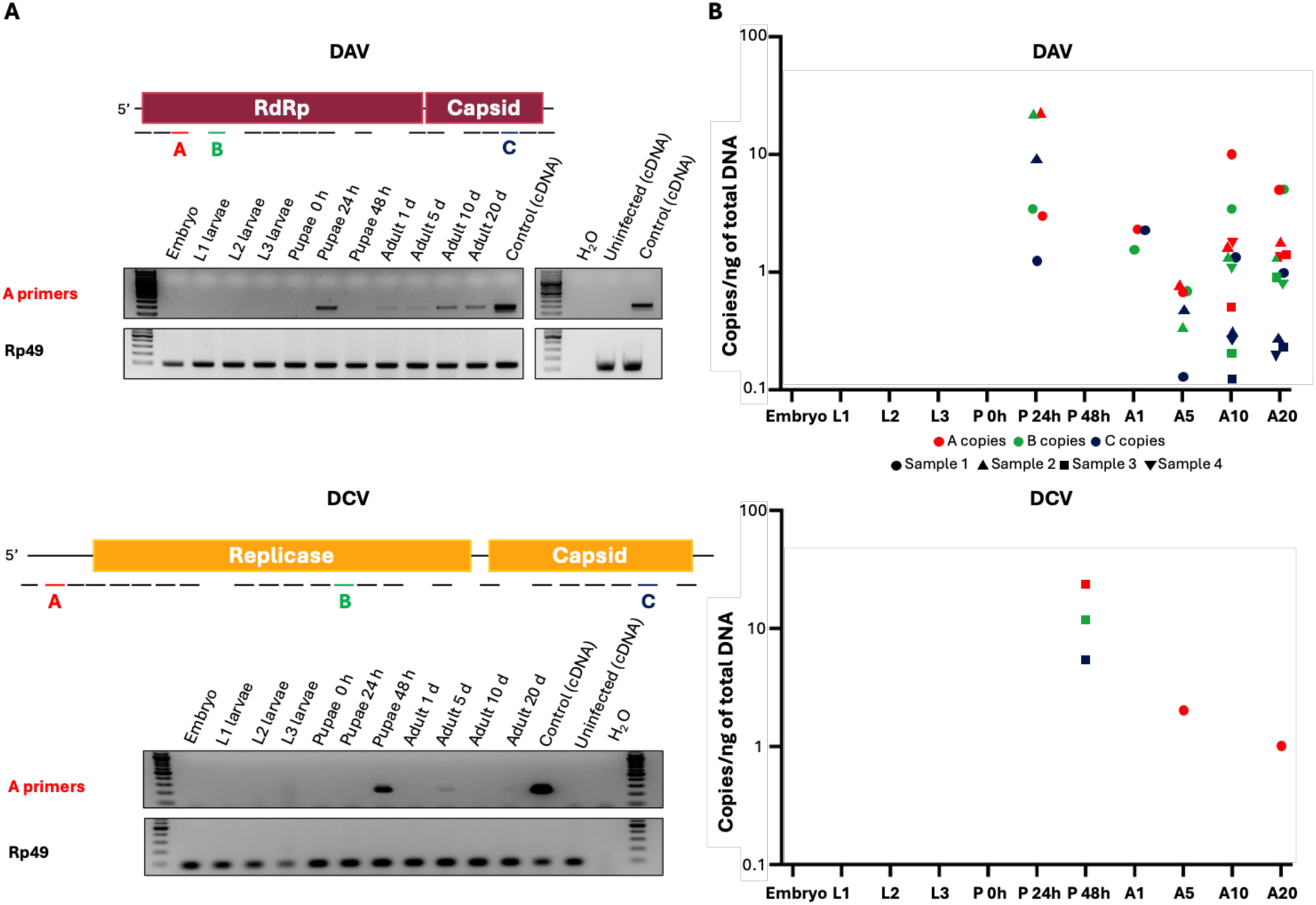
Viral DNA (vDNA) formation kinetics for DAV-and DCV-infected fly stocks. A) Schematic of genome-wide vDNA screening and fragment detection. Viral genomes were screened using ∼200 bp amplicons. Amplicons corresponding to detected vDNA are shown as black or colored lines below the viral genome they correspond to. PCR for detection of fragment A (highlighted in red in genome schematics) in the infected pools is shown. Each pool is combined DNA from 4 individuals at the same developmental stage. Control lanes included: infected sample cDNA (positive), uninfected fly cDNA (negative), and H₂O (no-template control). Rp49, a housekeeping gene was used as an amplification control. All gels used 100 bp ladders (Thermo Scientific 10588170) for reference. B) Quantitative analysis of selected vDNA fragments. Based on initial screening, three regions (A, B, and C) were quantified using qPCR with standard curves. Data represent vDNA copies normalized to total DNA content (copies/ng) from individual biological replicates (n=4 per stage), corresponding to the same samples pooled in panel A. Colors represent the targeted fragment of the viral genome, and shapes correspond to the individual sample in which it was detected.

We next quantified vDNA levels across developmental stages in the individuals that had been used to prepare the DNA pools tested by PCR. This was done by performing qPCR for the three DAV and DCV amplicons that were previously validated by Sanger sequencing (designated as primer sets A, B, and C in Fig. 3A). In DAV-infected insects, vDNA was first detected at 24 hours post-pupation—coinciding with peak RT activity— for all three amplicons, with levels reaching 22.35 copies/ng DNA (Figure 3B). vDNA levels subsequently fell below detection limits during late pupation but vDNA reemerged in adults beginning at eclosion (day 1), reaching 4.86 copies/ng DNA by 20 dpe (Figure 3B). Notably, we found that DAV replication levels were not correlated with vDNA levels (Supplementary Figure 6).

DCV showed more limited vDNA production, with vDNA first detected at 48 hours post-pupation at 24.12 copies/ng DNA. Unlike DAV, DCV vDNA detection in adults was sporadic, with only fragment A detected in single individuals at 5 and 20 dpe (2.05 copies/ng DNA and 1.03 copies/ng DNA, respectively), despite testing all three primer sets (Figure 3B). Although low-abundance DCV vDNA restricted robust statistical analysis in adults, the correlation observed across all vDNA-positive samples (Supplementary Figure 6) suggests a replication-linked vDNA dynamic distinct from DAV. No Nora virus-derived vDNA was detected at any developmental stage despite Nora virus inducing significant RT activity during pupation (Supplementary Figure 5) and maintaining active replication throughout development (Figure 1). All amplicons from Nora virus-infected samples were confirmed via Sanger sequencing to derive from non-specific host genomic sequences.

These results reveal fundamental differences in vDNA synthesis among RNA viruses: DAV and DCV generate transient vDNA during pupation then again during adulthood, and Nora virus produces no detectable vDNA despite active infection and high levels of RT activity at several stages.

### DAV vDNA sequences comprise both viral sequences and virus-host chimeras

Having identified pupation as the developmental stage where vDNA first appears and is at its highest levels, we next characterized the molecular nature of these vDNA forms. To determine whether vDNA represents viral sequences alone or chimeric molecules containing host genetic elements, we performed high-throughput sequencing on total DNA extracted from pools of 20 DAV-infected pupae at 24 hours post-pupation, the timepoint of peak vDNA detection.

Illumina double-stranded DNA (dsDNA) sequencing generated 544,145,817 paired-end reads, of which 168 pairs aligned to the DAV genome, distributed across its entire length (Figure 4A). Notably, only 6 of these 168 read pairs (3.6%) represented chimeric sequences containing both viral and host genomic elements. All chimeric junctions mapped to the central part of the viral genome, between nucleotide positions 2,000 and 4,000, primarily within the viral RNA-dependent RNA polymerase (RdRP) gene, with one junction in the capsid gene (Supplementary Figure 7). None of the non-viral sequences within the chimeric reads corresponded to a transposable element. The low recovery of viral sequences using a sequencing approach highlights the major challenges inherent in detecting vDNA using short-read technologies.

**Figure 4.**
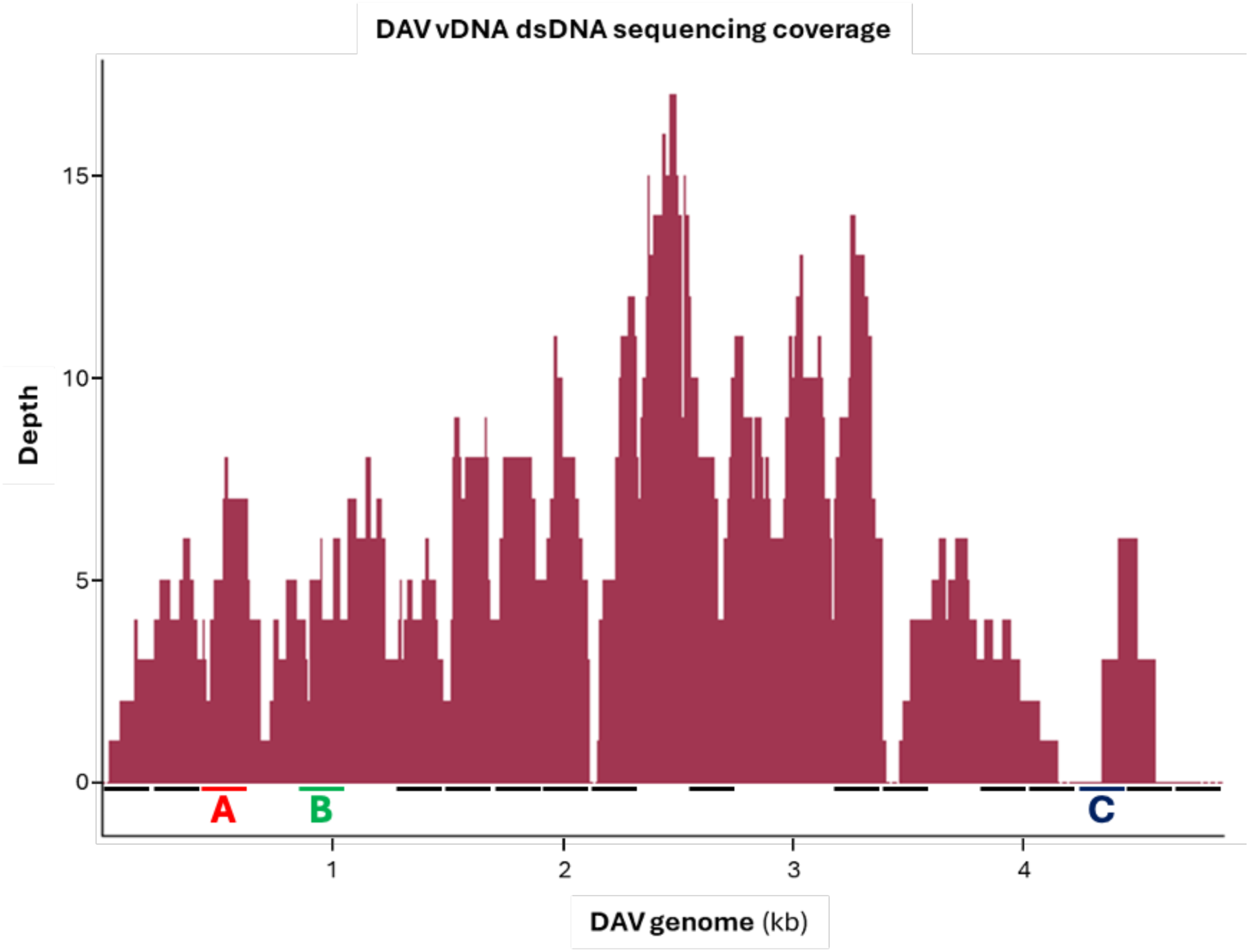
vDNA characterization in DAV-infected pupae at 24 h post-pupation. A) Depth profile (number of times a nucleotide is read during a sequencing run) of dsDNA paired-end sequencing reads (150 bp) mapping to the DAV genome (n=168 reads; complete or partial matches). The X-axis represents DAV genome position (kb). The positions of the A, B, and C fragments shown in Figure 3 are indicated.

### RNA virus infection modulates transposable element expression in adults

Transposable elements (TEs) are the source of endogenous RT activity in *Drosophila* and can contribute sequences to vDNA chimeras, as demonstrated for Flock House virus (Goic et al., 2013). Having observed that virus-induced RT activity peaks during pupation, we investigated whether viral infections also modulate TE expression in adults, where persistent infections are maintained. We analyzed RNA-seq datasets from adult *w^1118^* flies (1 and 12 dpe) that were uninfected or mono-infected with DAV, DCV, or Nora virus (Castelló-Sanjuán et al., 2025). These represent the same fly populations used throughout this study at timepoints common for all viruses, allowing us to connect adult TE dynamics with our developmental observations.

Transcriptional profiling revealed virus-specific modulation of TE expression that evolved from early to late infection. At 1 dpe, shortly after the pupal RT activity peak, DCV and Nora virus infections significantly upregulated long terminal repeat (LTR) retrotransposons, including DMCOPIA, DME487856, and TIRANT (Figure 5). By 12 dpe, when persistent infections are established, DAV infection induced broad upregulation across DNA, LINE, and LTR transposon families (Supplementary Figure 8), with LTR elements DMIS176, ACCORD, and DMCOPIA exhibiting the highest fold-changes (Figure 5). DCV infection consistently elevated DMCOPIA and DME487856 at both timepoints, while additionally inducing the LINE element DMRER2DM. Nora virus infection increased expression of DMCOPIA, DME487856, TIRANT, and DMRER2DM, and uniquely maintained continuous overexpression of the LTR element JUAN across both timepoints (Figure 5).

**Figure 5.**
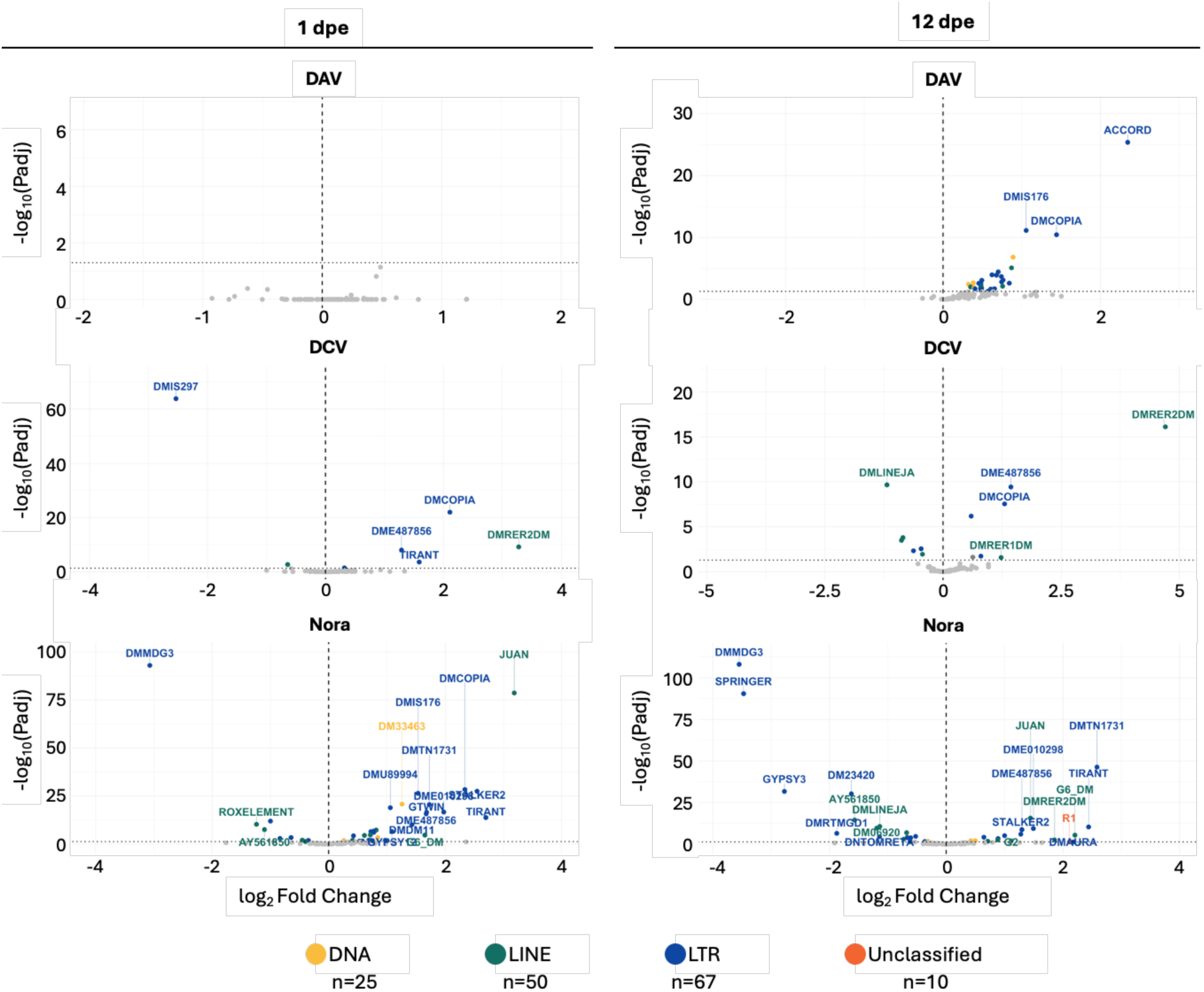
Differential expression analysis of transposable elements (TEs) in virus-infected Drosophila adults. Volcano plots showing differences in TE expression in mono-infected samples (DAV, DCV, or Nora virus) compared to uninfected controls at 1 day post-eclosion (dpe; left panels) and 12 dpe (right panels). Each point represents a TE from one of four orders: DNA, Long interspersed nuclear element (LINE), Long terminal repeats (LTR), or unclassified (total n=152: 25 DNA, 50 LINE, 67 LTR, 10 unclassified). Y-axes show-log₁₀(adjusted P-value), with horizontal dotted lines indicating the significance threshold (P_adj_ = 0.05). X-axes display log₂ fold change, where positive values indicate upregulation in the virus-infected samples and negative values indicate downregulation. Only TEs with |log₂FC| > 1 are labeled. Point colors represent TE classification: red (DNA), yellow (LINE), green (LTR), blue (unclassified).

Several upregulated TEs (DMCOPIA, DMRER2DM, and JUAN) match families previously identified in chimeric vDNA sequences generated during Flock House virus infection, with “Copia” elements producing most of the chimeras identified (Goic et al., 2013). However, TE upregulation seems to be dynamic. We observed upregulation of DMRER2DM (DCV and Nora virus), JUAN (Nora virus), and DMCOPIA (DAV and DCV), in a virus-specific manner. This overlap suggests that specific TE families may preferentially contribute to RT activity for vDNA synthesis. For instance, DMCOPIA expression is strongly induced during DAV and DCV infections, which produce detectable vDNA, while Nora virus infections show a distinct TE expression profile that may explain the absence of detectable Nora virus-derived vDNA despite the presence of Nora virus-induced RT activity.

These results suggest that viral infections established during early stages of development and pupation continue to shape the TE expression landscape in adults, potentially resulting in maintained RT activity that contributes to ongoing vDNA production and persistent infection.

### RNA virus infection reprograms pupal immune gene expression

So far, our results pointed towards pupation as a critical window for virus replication and RT activity. To investigate the underlying immune mechanisms during this window, we profiled the expression of key immune genes throughout pupal development. RNA from pupae with confirmed viral replication (Figure 1) was pooled by timepoint (0, 24, 48 hours post-pupation) and infection status (uninfected, DAV, DCV, or Nora virus). A total of 3 pools per developmental stage and infection were used. Gene expression was analyzed by RT-qPCR targeting components of major antiviral pathways. For each sample, gene expression values were normalized over a housekeeping gene (*Rp49*) in the same individual, using the ΔCt method. Expression values can be found in Supplementary File 1 and are summarized in Figure 6.

**Figure 6.**
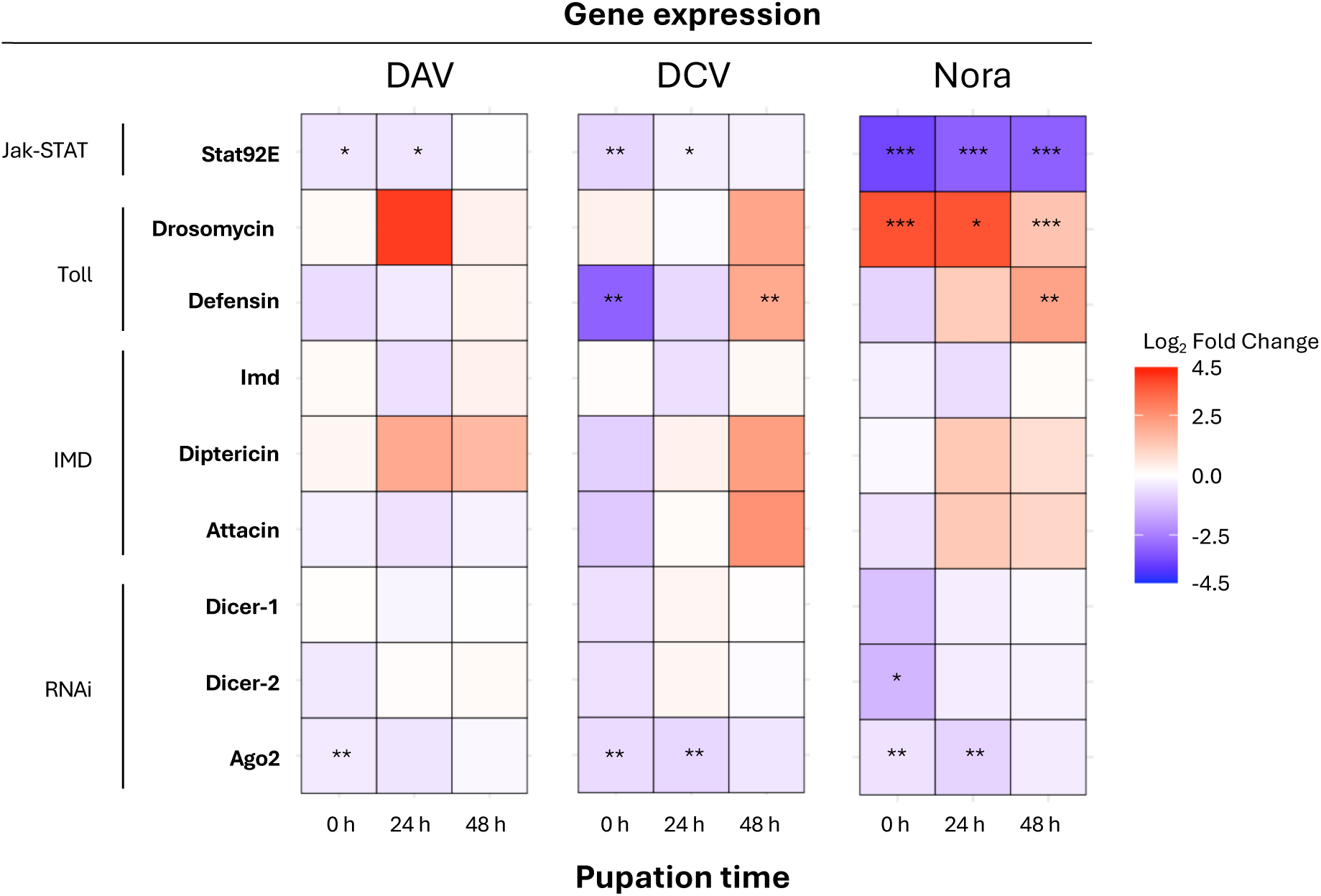
Immune landscape remodeling during persistent viral infection in pupae. A) Immune gene expression profiling in virus-infected pupae. RNA from three individual pupae (per time point and viral stock) with confirmed viral replication was pooled, and a total of 3 replicate pools were analyzed via RT-qPCR. For each sample, gene expression values were normalized to a housekeeping gene (Rp49) in the same individual using the ΔCt method. Expression values can be found in Supplementary File 1. Heatmaps display median log₂ fold changes relative to stage-matched uninfected controls. Genes are grouped by immune pathway (y-axis), while viral stocks (DAV, DCV, Nora virus) and pupation times are organized along the x-axis. Generalized linear models (GLM, Gaussian family) were used to compare infected versus uninfected samples for each gene-virus combination. Significance levels: *P < 0.05, **P < 0.01, ***P < 0.001 (nominal P-values).

All three viruses suppressed the RNAi pathway, the primary antiviral defense in insects, during pupation. Argonaute-2 (Ago2), which cleaves viral RNA targeted by siRNAs, was significantly downregulated at 0 h and 24 h post-pupation in all viral infections (Figure 6). Nora virus additionally suppressed Dicer-2 at 0 h, the enzyme that initiates RNAi by processing viral dsRNA into siRNAs, representing a more comprehensive inhibition of this antiviral pathway (Figure 6).

The Jak-STAT pathway, which coordinates systemic antiviral responses, was also suppressed by viral infection. Expression levels of Stat92E, a transcription factor that activates antiviral gene expression, decreased at 0 h and 24 h post-pupation across all three viral infections. At 48 h post-pupation, Stat92E expression remained suppressed during Nora virus infection, but was restored to normal levels during DAV and DCV infection (Figure 6).

Expression of antimicrobial peptides (AMPs), which are typically associated with antibacterial immunity but are increasingly recognized in antiviral defense, revealed virus-specific patterns. Drosomycin, an antifungal peptide regulated by the Toll pathway, was robustly upregulated throughout Nora virus infection (0 h, 24 h, and 48 h), but remained unchanged in DAV and DCV infections (Figure 6). Defensin, a broad-spectrum AMP, showed complex regulation: DCV infection caused early suppression (0h) of Defensin followed by late induction (48h), while Nora virus only induced expression of Defensin at 48 h (Figure 6).

These results reveal that viral infections converge on suppressing core antiviral pathways (RNAi and Jak-STAT) during early pupation, coinciding with peak viral replication and RT activity. However, each virus also induces distinct effector responses, with Nora virus showing the most extensive immune modulation by combining sustained Jak-STAT suppression, enhanced RNAi inhibition through both Dicer-2 and Ago2, and unique activation of the Toll/antimicrobial peptide response throughout pupal development.

## Discussion

Our study of infections with three phylogenetically distinct enteric RNA viruses across *D. melanogaster* development demonstrates that pupal metamorphosis is a key node when viral replication, RT activity, vDNA synthesis and immune regulation converge, potentially defining the fate of an infection (lethality vs. persistence vs. clearance). This finding aligns with emerging evidence that insect metamorphosis acts as an important developmental window that influences antiviral responses (Wang et al., 2022).

The significance of metamorphic transitions in shaping host-virus interactions extends across examples from diverse insect taxa. In Lepidoptera, *B. mori* shows progressively less susceptibility to Bombyx mori nuclear polyhedrosis virus as pupation progresses, indicating a refinement of antiviral mechanisms during metamorphosis (Mikhailov et al., 1992). In Diptera, *A. aegypt*i pupal-specific factors exert antiviral influence. The pupal cuticle protein directly suppresses dengue virus replication when expressed in mosquito cells (Huang et al., 2023). In Hymenoptera, during *Apis mellifera* pupation, pupae show significant changes in gene expression and DNA methylation in response to viral infection, such as Israeli Acute Paralysis Virus, indicating that the pupal stage is both a vulnerable period and a time of active immune regulation (Li-Byarlay et al., 2020). Collectively, despite the variety of models and implications, these studies highlight that the metamorphic transition from larval to adult stages harbors pivotal molecular and physiological factors that influence infection outcomes. While embryonic infection (e.g., vertical transmission in Nora virus) can modulate initial viral acquisition, pupation represents a universal developmental reprogramming event that actively reshapes infection trajectories through host immune remodeling and tissue reorganization, irrespective of transmission mode. Although the importance of this developmental stage across insects is recognized, the mechanistic processes underlying how different RNA viruses interact with pupal development to establish persistence have remained largely unexplored.

Our results reveal that viral RNA dynamics follow virus-specific trajectories throughout fly development. DAV-and DCV-infected populations show peaks in viral RNA levels at 24 hours post-pupation. Conversely, Nora virus-infected samples maintain stable viral RNA levels throughout development, indicating distinct regulatory mechanisms. Despite these different patterns, all three viruses show peaks in prevalence and replication levels during pupation that coincide with increased endogenous RT activity. Since these viruses do not encode reverse transcriptases, any changes in RT activity stem from the viruses’ impact on endogenous retrotransposon activity. It has been shown that modulation of TEs affects developmental control, morphogenesis, central nervous system function, immune responses, homeostasis, infection resistance, and survival (Balakireva et al., 2024; Ullastres et al., 2021; Wang et al., 2022) and that viral infections modulate the expression levels of host TEs. In *A. aegypti*, arboviral infections regulate TE expression in a virus-specific manner (Garambois et al., 2024). In *Drosophila (melanogaster and simulans)* acute or chronic viral infections impact TE activity, and thus the tempo of genetic diversification (Roy et al., 2020). Additionally, in *D. melanogaster* the retrotransposon mdg4 (Gypsy) shapes innate immunity during metamorphosis through the dSTING-NF-κB/Relish signaling pathway, priming antiviral defenses in adulthood (Wang et al., 2022). This coordination suggests that pupal physiology creates conditions that promote both viral replication and retrotransposon activation. This reflects findings in mammals, where host factors that limit viral infections (e.g., APOBEC3, TREX1, SAMHD1) similarly suppress retrotransposons, highlighting a fundamental mechanistic link between antiviral immunity and TE control across species (Goodier, 2016; Goodier et al., 2015; Wichroski et al., 2006).

The molecular signature of virus-TE interactions manifests through vDNA synthesis, which occurs in virus-specific patterns. For instance, in *Aedes* mosquitos infected with Semliki Forest Virus, vDNA molecules are present as episomes and can be found as chimeras with several LTR retrotransposon sequences (Rodriguez-Andres et al., 2024). However, the abundance and types of TEs in mosquitoes are different from those found in *Drosophila*, where sequencing of vDNA:TE chimeras has thus far only been successful for FHV infection and only during *in vitro* infections (Goic et al., 2013; Poirier et al., 2018). While we successfully quantified and sequenced vDNA from *in vivo* infections for this first time in this study, our findings show that detection of vDNA with current sequencing methods remains challenging, as we obtained very few vDNA-derived reads. We tested several strategies to enrich vDNA prior to sequencing (probe-capture, rolling-circle amplification using Phi29, and PCR-based strategies), but these ultimately did not increase vDNA yield (data not shown), highlighting inherent limitations in detecting these molecules. These findings indicate that future efforts should prioritize alternative enrichment methods and long-read technologies to overcome resolution challenges and improve vDNA sequencing. These technical challenges in vDNA detection could arise from secondary structures within vDNA molecules, from the potential existence of vDNA as RNA-DNA hybrids, from localization within specific cellular compartments or tissues that are refractory to DNA extraction methods, or from association with proteins that hinder sequencing accessibility. Additionally, vDNA molecules often contain extensive repetitive sequences, which inherently pose difficulties for quantification, sequencing, and subsequent bioinformatic analysis (Delahaye & Nicolas, 2021; Treangen & Salzberg, 2011; Zattera & Bruschi, 2022).

Transgenerationally-inherited vDNA has been detected in uninfected adult mosquitoes and in the uninfected progeny of infected *D. melanogaster* flies (Mondotte et al., 2018, 2020; Rodriguez-Andres et al., 2024). We did not detect vDNA in the embryos of infected adults in this study. However, we did detect short viral RNA fragments in some embryos from DAV-and DCV-infected parents. These observations raise the intriguing possibility that viral RNA fragments may be subjected to transgenerational transmission and may potentially act as a template for vDNA synthesis upon recognition by retrotransposons as part of a mechanism for heritable viral immunity.

Virus-specific TE activation patterns provide insight on differential vDNA formation. Our analysis reveals shared upregulation of the LTR retrotransposon DMCOPIA in DAV and DCV infections, consistent with prior vDNA sequencing of FHV-infected cells (Poirier et al., 2018). Similar Copia-like LTR elements have been identified in vDNA chimeras in *A. aegypti* mosquitoes infected with positive-sense RNA arboviruses (Rodriguez-Andres et al., 2024). Additionally, DCV and Nora virus infections show upregulation of the LINE TE DMRER2DM, previously linked to FHV vDNA (Poirier et al., 2018), while Nora virus infection also upregulates JUAN, another LINE TE associated with chimeric FHV vDNA (Poirier et al., 2018). These findings indicate virus-specific TE interactions, likely driven by compatible virus-retrotransposon partnerships, which may account for the absence of Nora virus-derived vDNA.

A previous study found that hemocytes are the primary cell type in which synthesis of vDNA occurs (Tassetto et al., 2017). We observed that vDNA synthesis begins and virus-induced RT activity peaks in pupation, the same developmental window in which the first fully differentiated hemocytes are released into the hemolymph through a process that requires Jak-STAT activity (Banerjee et al., 2019, Trivedi & Starz-Gaiano, 2018). Our observation that viral infection downregulates Stat92E during pupation thus raises the intriguing possibility that viruses may target Jak-STAT signaling to limit vDNA synthesis by disrupting hemocyte differentiation during pupation.

Our findings establish pupal metamorphosis as a critical developmental window during which viral infections interact with the immune mechanisms of the developing organism, potentially contributing to the establishment of persistent infections in adults. During this stage, endogenous retrotransposon activity, immune pathway modulation, and vDNA synthesis capacity intersect with viral replication dynamics. Different viruses navigate these host changes through distinct mechanisms and achieve persistence. Future studies will address whether and how all these physiological observations come together to coordinate the establishment and maintenance of viral persistent infection through developmental constraints.

## Materials and methods

### Fly stocks

For all experiments, we used *Wolbachia*-free *w^1118^* flies maintained on a standard cornmeal diet. This diet was prepared by combining 440 g inactive dry yeast, 440 g corn meal, and 60 g agar in 6L osmotic water. The mixture was autoclaved and after cooling, 150 ml moldex solution (20% methylhydroxybenzoate) and 29 ml propionic acid were added as preservatives. All flies were kept at 25°C under a 12:12 light:dark cycle. All fly lines were cleaned of possible persistent infections (viruses and *Wolbachia*) as described previously (Merkling & van Rij, 2015).

### Persistent infected fly lines

Persistently infected *w^1118^ Drosophila melanogaster* lines were established following the methodology described by Nigg and colleagues in 2022, aligning with the stocks utilized in their subsequent research (Nigg et al., 2024). Four RNA virus persistent infections were introduced to the *w^1118^* flies using specialized techniques. For Drosophila C virus (DCV), a persistently infected bw1;st1 Ago3t3/TM6B;Tb+ strain (Bloomington #28270) served as the source to transfer DCV to naïve *w^1118^* flies through environmental exposure. The infected donor flies were kept in a new vial for three days; thereafter, the *w^1118^* flies were allowed to be exposed to the contaminated environment for an additional three days. The F0 generation was then transferred to a fresh vial, where their F1 progeny were confirmed to be persistently infected.

For Drosophila A virus (DAV), an Australian isolate (DAVHD from the van Rij lab) was injected into adult *w^1118^* flies at a volume of 50 nl per fly. Following a similar environmental transfer procedure, the injected flies contaminated the vial for three days before naïve *w^1118^* flies were exposed. Persistently infected F1 progeny were isolated and maintained accordingly. The Bloomfield virus was detected in a Dipt-GFP stock (BDSC Cat#55709). A homogenate from these flies was filtered (0.22 µm) and subsequently injected into adult *w^1118^* flies. After injecting the F0 flies, they were removed after three days, while F1 and F2 progeny underwent successive generational passages (5–9 days per vial) to establish persistence in F3 adults, which was confirmed via RNA sequencing.

The Nora virus-persistent line originated from a *w^1118^* line infected with Nora virus, which was kindly provided by Dr. Stefan Ameres. Upon receiving the sample, we performed repeated backcrossing of the infected *w^1118^* flies with the *w^1118^* line maintained in our lab. We subsequently verified that the genetically backcrossed flies continued to show persistent infection with Nora virus through RNA sequencing. All stock lines were preserved at 25°C under a 12:12 light:dark cycle. The presence or absence of persistent infections was assessed using RT-PCR with specific primers for Nora virus, DCV, and Drosophila A virus (Supplementary Table 2). Additionally, total RNA from the persistently infected adults was extracted and sequenced. The resultant reads were aligned to a comprehensive database of known Drosophila viruses (https://obbard.bio.ed.ac.uk/data.html), confirming the exclusive presence of the targeted virus (Nigg et al., 2022, 2024).

### Developmental stage sample collection

From the uninfected and infected fly stocks, 20 males and 20 females were placed in clean fly food vials to lay eggs for 16 hours. After 16 hours, the parental generation was removed, and the offspring were left in the tube to develop. To collect the embryo and larval samples, a 20% sucrose solution was dispensed onto the food, and the food was gently disrupted with a brush (adapted from Nichols et al., 2012). Tubes with the sucrose solution were left for 20 minutes to allow the disrupted food particles to sediment and the larvae and embryos to float. Then, with the use of a 1000 μl pipette with a cut tip, embryos and larvae were collected onto a fine mesh (30 μm, Flystuff). This mesh was placed over a tube, and the liquid was discarded. Embryos and larvae were rinsed with MilliQ H₂O to prevent any downstream effects of food or sucrose presence. To further ensure the detection of nucleic acids coming from within the live embryo host, embryos were bleached with a 50% bleach solution (Merkling & Van Rij, 2015). Pupae were collected using Drosophila sorting brushes dipped in MilliQ H₂O. The use of water detached the pupae from the tube’s surface without damaging them. Adults were anesthetized with CO₂ and collected using Drosophila sorting brushes over a CO₂ pad (Flystuff® Flypad).

### RT activity protocol

RT activity was measured in whole fly extracts using a protocol previously adapted from Pyra et al., 1994 and Wu et al., 2021 (Bergman et al., 2025). Six to nine pools of 15 to 20 individuals of the same developmental stage, from different uninfected or persistently infected flies, were collected. For the adult samples, individual samples were collected and prior to use, an aliquot underwent RNA extraction to pool only those infected. For all the developmental stages pools or individuals were collected into 100 μL of CHAPS lysis buffer (Goic et al., 2013), containing 10 mM Tris-HCl pH 7.5 (Invitrogen, AM9855G), 400 mM NaCl (Invitrogen, AM9759), 1 mM MgCl2 (Invitrogen, AM9530G), 1 mM EGTA (Sigma-Aldrich, E4378), 0.5% CHAPS (Thermo Fisher Scientific, 28300), 10% glycerol (Sigma-Aldrich, J61059-AP), 1 mM DTT (Invitrogen, Y00147), and 1X cOmplete EDTA-free protease inhibitor cocktail (Roche, 11873580001).

The tissues were homogenized for 30 seconds at 5500 rpm in a Precellys homogenizer (Bertin Technologies) and stored at-80°C. Subsequently, the samples were clarified by centrifugation for 10 minutes at 16,000 g to remove tissue debris. The resulting supernatant was collected into a new tube. Protein content in the supernatant was measured using Pierce Detergent Compatible Bradford Assay kit (Thermo Fisher Scientific, 23246) according to the manufacturer’s instructions. All samples were diluted to the working concentration of 1 μg/μl. Prior to the RT reaction, 30 pmol of MS2 reverse primer (Primer_1, Supplementary Table 1) with 0.4 mM dNTPs (Thermo Fisher Scientific, R1121) were allowed to anneal to 100 ng of MS2 RNA (Roche, 10165948001) in a 12.5 μL annealing reaction at 65°C for 5 minutes. The annealing reactions were used in subsequent 25-μL RT reactions, containing 30 pmol MS2 reverse primer, 100 ng MS2 RNA, 0.2 mM dNTPs, 1 mM DTT (Invitrogen, Y00147), 10 mM Tris-HCl pH 7.5 (Invitrogen, AM9855G), 1 mM KCl (Invitrogen, AM9640G), 0.14 mM MnCl2 (Sigma-Aldrich, M1787), 0.02% Triton X-100 (Sigma-Aldrich, 93433), 40 units of RNase OUT (Invitrogen, 10777-019) and 5 μL of protein sample. The RT reaction was allowed to progress for 1 hour at 25°C, after which it was stopped by heat-inactivation at 70°C for 15 minutes. Heat-inactivated samples (45 minutes of incubation at 98°C), CHAPS buffer alone, and no-template reactions served as negative controls. A positive reference control of 1 unit of SuperScript II (Invitrogen, 18064022) was used for relative quantification of RT activity.

Generated MS2 cDNA was quantified with three technical replicates through quantitative PCR (qPCR) in a 10 μL reaction containing 3 pmol of MS2 forward primer and reverse primers (Primers 1-2, Supplementary Table 1), and 1 μL of RT reaction (template) using Luminaris Color HiGreen qPCR Master Mix, low ROX ThermoFisher, K0374). A calibrator sample of known MS2 cDNA was added to all the plates. Analysis of RT activity in the samples was done relative to the amount of cDNA of the SuperScript II wells.

RT activity data were analyzed in R (v4.3.2). Shapiro-Wilk tests confirmed non-normality (all W < 0.97, *P* < 0.05). Primary analysis used generalized linear mixed models (GLMM; lme4 package) with Gamma distribution (log link function), including condition (infected/uninfected), developmental stage, their interaction as fixed effects, and experiment as random intercept. Stage-specific infected vs. uninfected comparisons were conducted via Tukey-adjusted pairwise contrasts of estimated marginal means (emmeans package). Model diagnostics included residual plots to verify distributional assumptions. Analyses were performed separately for DCV and Nora viruses (α = 0.05).

### RNA extraction

For all the developmental stages, individual flies were collected in 100 μl of DPBS (Gibco, 14190144) and homogenized using a pestle. For those individual female flies subsequently used in the RT activity assay, individual flies were homogenized in CHAPS buffer (Goic et al., 2013), and processed as detailed in the RT activity protocol methods section. For all samples, an aliquot of 50ul of the homogenate was used for RNA extraction using TRIzol reagent (Invitrogen, 15596026). RNA was extracted following manufacturer’s instructions, supplemented with GlycoBlue (Invitrogen, AM9516) during the precipitation process. RNA concentrations were measured using the Qubit RNA BR Assay Kit (Invitrogen, Q10211).

### DNA extraction

For all the developmental stages, individual flies were collected in 100 μl of DPBS (Gibco, 14190144) and homogenized using a pestle. For all samples, an aliquot of 50ul of the homogenate was used for DNA extraction using Phenol:chloroform:isoamyl alcohol (Sigma-Aldrich, P3803) following the manufacturer’s instructions, supplemented with GlycoBlue (Invitrogen, AM9516) during the precipitation process. DNA concentrations were measured using the Qubit DNA dsDNA Assay kit (Invitrogen, Q32851).

### Standard preparation for quantification of viral replication and vDNA levels

qPCR amplicons were generated by RT-PCR from RNA extracted from persistently infected flies. Briefly, RNA was reverse transcribed using a T7-tagged reverse virus primer with SuperScript II reverse transcriptase (Sigma, 18064014), following the manufacturer’s instructions (Primers 4,7,9, Supplementary Table 1). The resulting cDNA served as a template for PCR amplification with DreamTaq DNA polymerase (Thermo Fisher Scientific, K1081), also according to the manufacturer’s protocol. Primers included the T7 promoter and a virus-specific forward primer (Primers 5,6,8,10, Supplementary Table 1). PCR products were purified using the Macherey-Nagel PCR purification kit (740609) as per the manufacturer’s instructions, and their identities were confirmed through Sanger sequencing. The purified PCR products were cloned into the pCR4-TOPO vector utilizing the TOPO TA cloning kit (Thermo Fisher, K4575J10) according to the manufacturer’s instructions. Clones containing the insert in the reverse primer orientation under the control of the T7 promoter were used as templates for in vitro transcription of negative sense RNAs, employing the MEGAscript T7 in vitro transcription kit (Invitrogen, AMB13345), following the manufacturer’s guidelines. The transcribed RNAs were further treated with DNase I (Roche, 04716728001) to remove residual DNA, according to the manufacturer’s instructions, and purified via phenol:chloroform extraction followed by ethanol precipitation. The concentration of the purified RNAs was measured using Qubit BR (Thermo Fisher, Q10211), and these concentrations were used to estimate RNA copy numbers per milliliter based on the respective RNA sizes.

### RT-qPCR for quantification of viral load and replication levels

Reverse transcription was performed using random primers or strand specific primers (Primers 5,7,9, Supplementary Table 1) and Maxima H Minus Reverse Transcriptase (Thermo Scientific, EP0751) following manufacturer’s instructions. The resulting cDNA was diluted 1:10 with MiliQ water. The qPCR was performed using Luminaris Color HiGreen qPCR Master Mix (Thermo Scientific, K0374). The reactions were done in triplicates of 10 ul each and using the corresponding set of primers for each virus (Primers 5,6,8,10,11-18, Supplementary Table 1). The cycling conditions were as follows: 2 minutes at 50°C, 10 minutes at 95°C, followed by 40 cycles of 15 seconds at 95°C and 60 seconds at 60°C. A standard melt curve analysis was performed after the cycling protocol. For the analysis of the results, the threshold for viral RNA detection was set to Ct 35 (i.e. if a sample had Ct higher than 35 it was considered non infected).

For the analysis of the viral load, the Ct values of the virus for a sample were normalized against the Ct value of the housekeeping gene *Rp49* for that sample (ΔCt). The 2^-ΔCt^ values were calculated per sample and their log10 plotted. For the analysis of viral replication, previously an RNA standard curve was produced ranging from 10^8^ to 10^1^ molecules per µl. RT was performed in both the samples as previously described, using strand-specific primers to target the negative RNA strand. The obtained Ct values from the samples were used in the standard curve equation to find the number of negative strand viral RNA molecules in the tested samples.

Statistical analyses were performed in R (v4.3.2). Viral load data (DAV, DCV, Nora) were log₁₀-transformed to achieve normality, while replication data (inherently log-scaled) were analyzed untransformed. Normality was confirmed via Shapiro-Wilk tests. Linear mixed-effects models (lme4 package) included developmental stage as a fixed effect and experiment as a random intercept. Pairwise comparisons between stages were conducted using Tukey-adjusted contrasts via estimated marginal means (emmeans package). Significance (α = 0.05) is denoted in figures using compact letter displays (multcompView package).

### Primer design for vDNA and RT-PCR

Primers targeting DCV, DAV, and Nora virus were designed using Primalscheme (primalscheme.com). To ensure viral specificity and eliminate amplification of *Drosophila* host DNA, all primers underwent rigorous in silico validation. First, local BLAST analysis against the *Drosophila melanogaster* genome (release r6.56) identified potential off-target binding sites. We used the default values for short BLAST settings. Primer pairs were subsequently discarded if both primers aligned to the same chromosomal locus within a 400 bp interval, preventing nonspecific amplification of host genomic fragments.

### PCR for vDNA

Previously designed primers were sorted and ordered in plate format. Genomic DNA was extracted from four independent pools of infected stocks (DAV, DCV, Nora virus), each consisting of 10 adult female flies of unspecified ages. As a positive control, cDNA pooled from RT-qPCR–confirmed infected samples was diluted and used. Additionally, cDNA derived from RNA of a pool of 10 uninfected females served as the negative control, along with the H₂O negative control. PCR amplification was performed with DreamTaq DNA polymerase (Thermo Scientific, K1081) following the manufacturer’s conditions, using 35 cycles with a consistent annealing temperature of 60°C. Results were visualized on a 1.5% agarose gel. Three primer pairs were selected, targeting three different regions on the viral genome (A, B, and C; Primers 19-30, Supplementary Table 1) as well as an endogenous *Rp49* control (Primers 31,32, Supplementary Table 1). The PCR amplicon was Sanger sequenced and confirmed to be virus in origin. Following, PCRs were repeated on pooled DNA coming from four individual samples in which viral replication was confirmed.

### qPCR for vDNA

Previously extracted DNA from individual samples underwent qPCR using qPCR primers (Primers 31-44, Supplementary Table 1) targeting the A, B, and C regions used to follow vDNA formation by PCR, as well as an endogenous *Rp49* control. vDNA standards were used from 10^8^ to 10^1^ copies/reaction. For all reactions, uninfected and H_2_O controls were added. The qPCR was performed using Luminaris Color HiGreen qPCR Master Mix (Thermo Scientific, K0374). The reactions were done in duplicates of 10 ul each and using the corresponding set of primers for each virus (Supplementary Table 1). The cycling conditions were as follows: 2 minutes at 50°C, 10 minutes at 95°C, followed by 40 cycles of 15 seconds at 95°C and 60 seconds at 60°C. A standard melt curve analysis was performed after the cycling protocol. For the analysis, the standard curve values were used and the number of copies normalized over the input DNA.

### vDNA sequencing

Genomic DNA was isolated from pools of 20 DAV-infected pupae at 24 hours post-pupation using phenol:chloroform extraction. For double-stranded DNA sequencing, 500 ng of extracted DNA was fragmented by sonication (Covaris) and processed with the NEBNext Ultra II DNA Library Prep Kit (New England Biolabs, E7645).

Following this, bioinformatic processing began with quality assessment of raw reads using FastQC, followed by adapter trimming and quality filtering with Cutadapt. Processed reads were aligned to reference genomes using Bowtie2 under --sensitive and--end-to-end parameters, generating SAM files that were converted to sorted BAM files using SAMtools (view, sort, and index commands). Read depth profiles were calculated with SAMtools depth. For chimera detection, reads were converted to FASTA format and used to construct a BLAST database via makeblastdb (NCBI BLAST+). Putative chimeric sequences were identified through blastn alignment against viral genomes with an E-value threshold of 1×10⁻¹⁰.

### Pupa immunity

To analyze the expression levels of multiple immune-related genes, RT-qPCR was conducted on RNA extracted from infected pupae. RNA samples from three individual pupae, either uninfected or mono-infected with DAV, DCV, or Nora virus, collected at 0, 24, and 48 hours post-pupation, were pooled. cDNA synthesis was carried out using SuperScript II reverse transcriptase and random hexamer primers. qPCR was performed with gene-specific primers (Primers 45-64, Supplementary Table 1). All reactions included the housekeeping gene *Rp49* as an internal control for normalization (Primers 31, 32, Supplementary Table 1). Gene expression data were analyzed in R (v4.3.2). For each target gene and virus (DAV, DCV, Nora), generalized linear models (GLM) with Gaussian distribution and identity link function compared expression between infected and uninfected groups. Model specification: expression ∼ Stock. To address potential skewness, sensitivity analyses using log-transformed data yielded identical significance results. Effect estimates represent mean expression differences between infected and uninfected groups. *P*-values are nominal and unadjusted for multiple comparisons. All analyses employed two-tailed tests with an α level of 0.05.

### Transposable element analysis of available datasets

RNA-seq data were processed with the rnaflow pipeline (*hub / RNAflow · GitLab*, 2024). Briefly, reads were aligned to the D. melanogaster reference genome, with genome sequence, gene annotations, and TE consensus sequences obtained from FlyBase (Öztürk-Çolak et al., 2024), release 2023_02. Alignment was performed using STAR v2.7.11b (Dobin et al., 2013) in two-pass mode with options “-- outFilterMismatchNoverLmax 0.05 --outFilterMultimapNmax 50 --seedSearchStartLmax 20“. Reads were counted with TEtranscripts v2.2.3 (Jin et al., 2015) in “--mode multi”. Differential expression was analyzed on a combined gene and TE count table using DESeq2 v1.40.2 (Love et al., 2014), excluding genes and TE families with a total raw count below 10 across all libraries. Gene set enrichment was conducted using the DESeq2 results with the R package fgsea v1.26.0 (Korotkevich et al., 2016), grouping TE families by order. All visualizations were created with ggplot2 v3.5.1 (Hadley Wickham, 2016).

## Author contributions

M.C.-S., R.G., J.C.N., and M.-C.S designed the experiments; M.C.-S., R.G., A. B., H.B., L. F., and J.C.N. performed the experiments; M.C.-S., R.G., A.B., L. F., J.C.N., and M.-C.S analyzed the data; M.C.-S., and R.G., wrote the first draft with inputs from L. F., J.C.N., and M.-C.S. All authors reviewed and approved the final version of the manuscript.

## Acknowledgements

We thank all members of the Saleh lab and Louis Lambrechts for their insightful discussions. We especially thank Gael Cristofari, whose feedback and inputs have been crucial for this work. We thank Cassandra Koh for her comments on the manuscript. We also acknowledge the use of Biorender.com for the fly’s illustrations.

## Supplementary figures

**Supplementary Figure 1.**
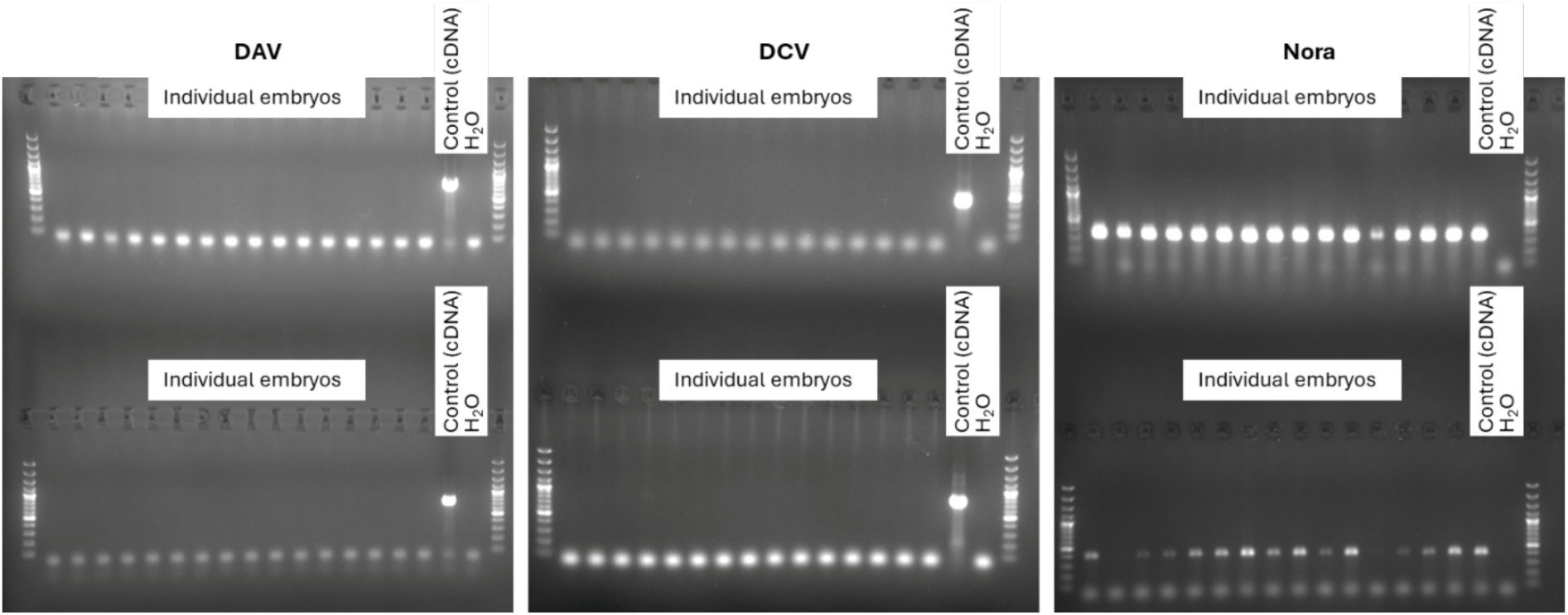
PCR results of viral genome amplification targeting 2 different regions. For all reactions, PCRs were run on cDNA coming from individual RNA samples that underwent reverse transcription with random primers. The cDNA of an infected adult sample serves as a positive control. H_2_O serves as negative control of the reaction.

**Supplementary Figure 2.**
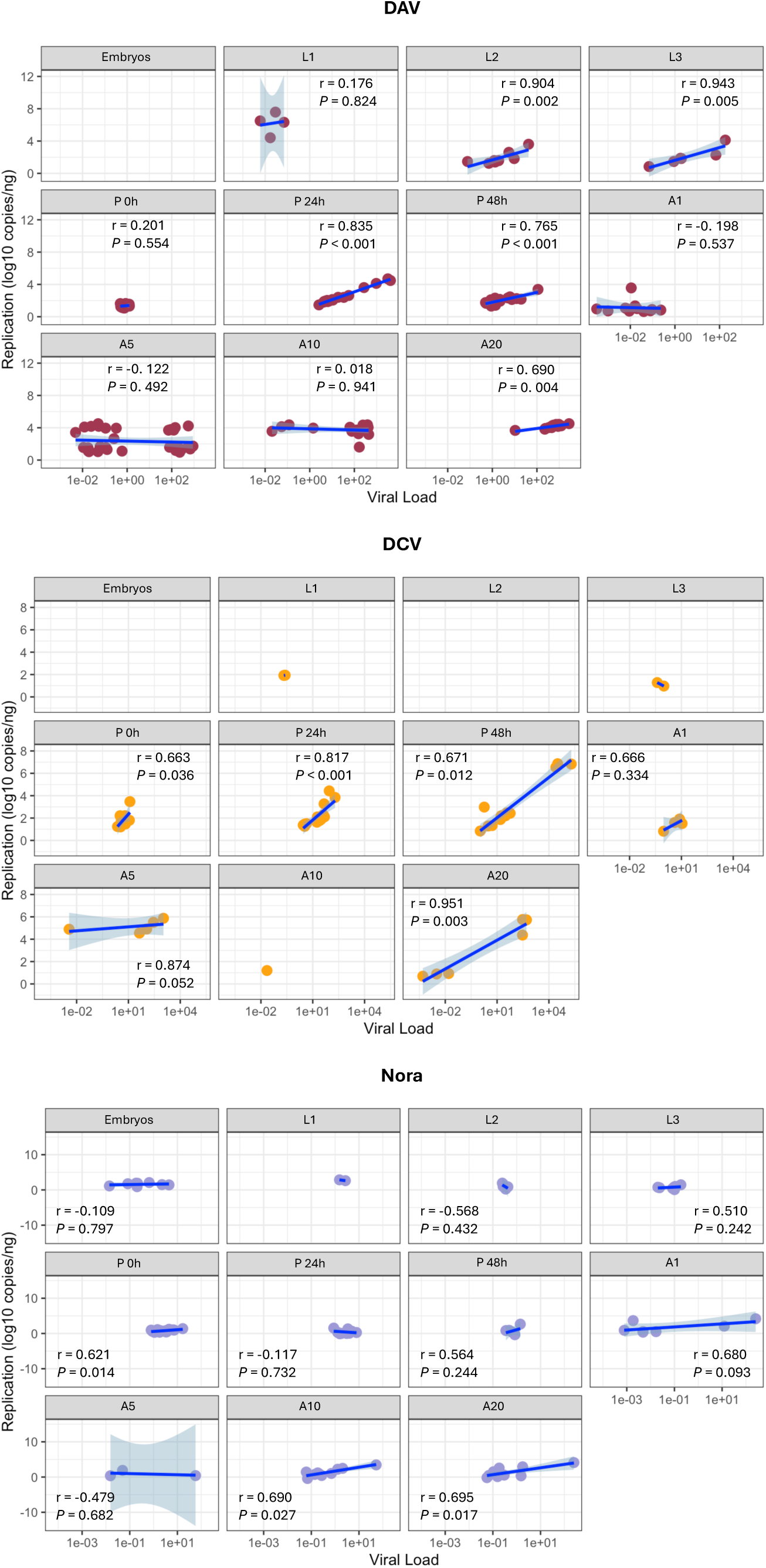
Replication vs. viral load in the infected samples throughout fly development. Results from Figure 1. For each individual sample, viral replication (Y axis) and viral load (X axis) are plotted. Samples are grouped per developmental stage and virus stock. Pearson Correlation values (r and P) are plotted in each graph.

**Supplementary Figure 3.**
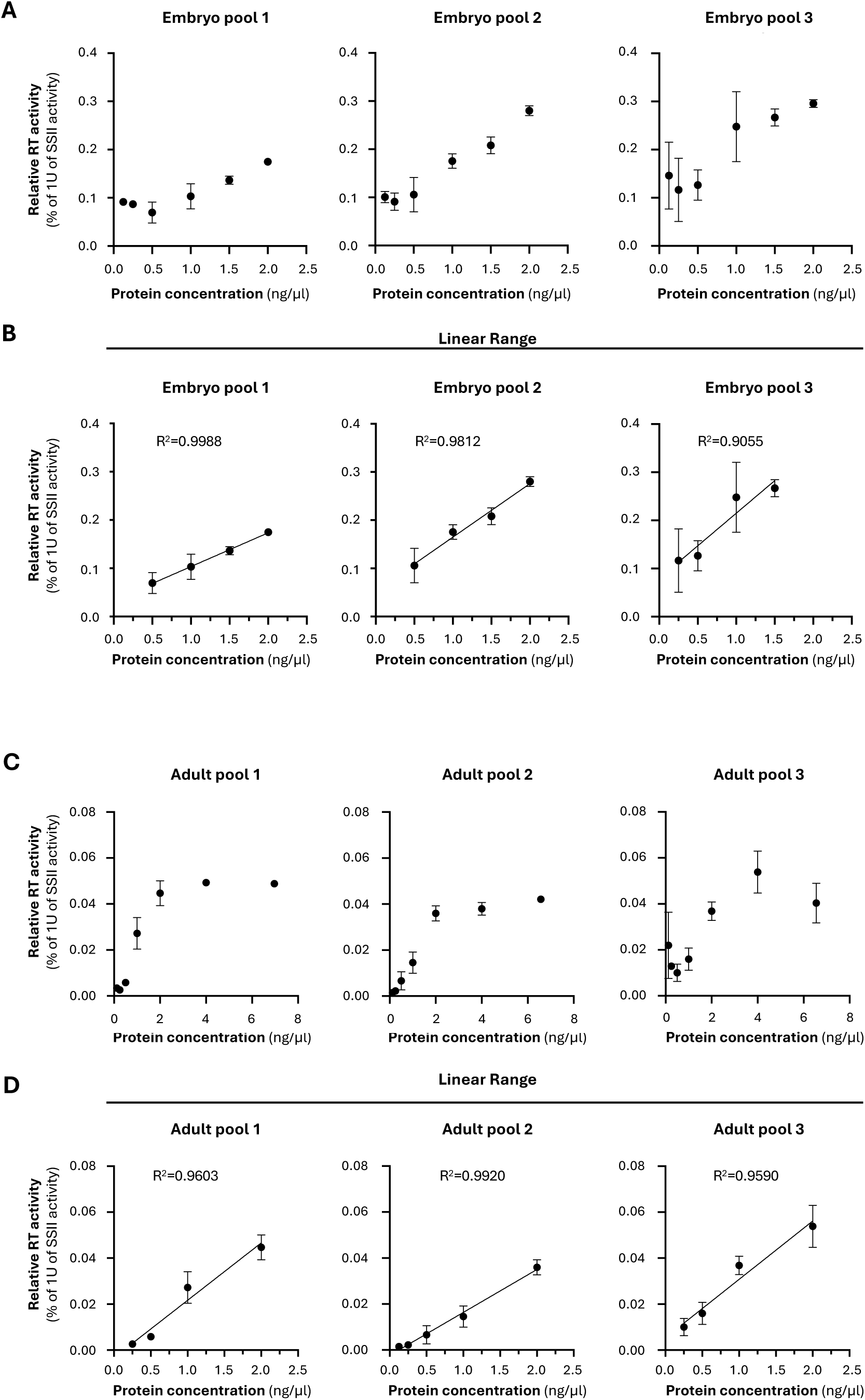
Linearity of the RT assay depending on the concentration of total fly protein.

**Supplementary Figure 4.**
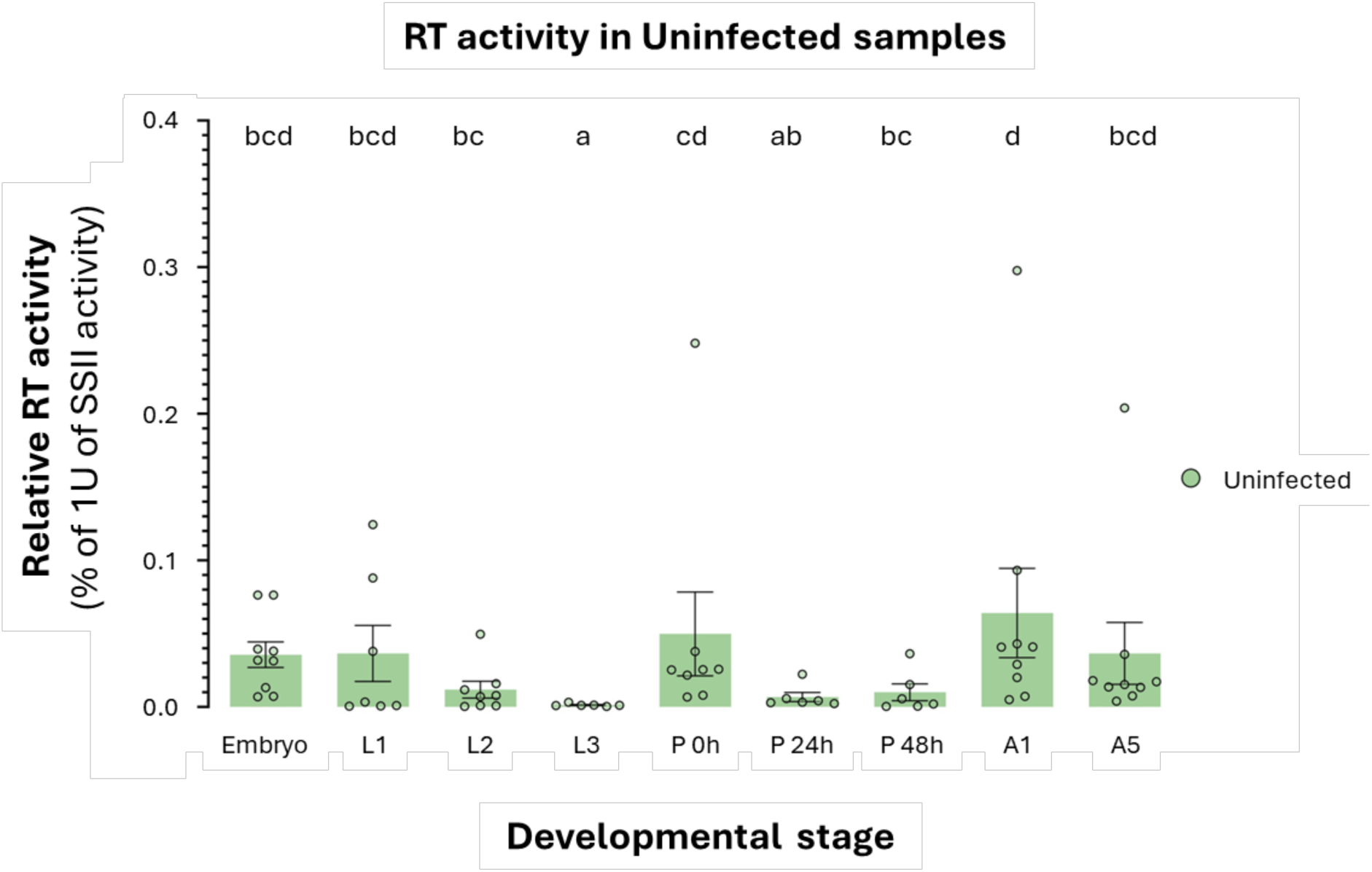
Relative RT activity of the uninfected samples. Uninfected samples were collected throughout development and processed to measure endogenous RT activity via RT-qPCR of an exogenous MS2 RNA. Y axis shows relative RT activity of the tested samples to that of 1 unit of SSII. X axis depicts developmental stages. RT activity in each developmental stage were compared to each other using generalized linear mixed models (GLMM, Gamma distribution) with experiment as random effect (P < 0.05, Tukey-adjusted pairwise comparisons).

**Supplementary Figure 5.**
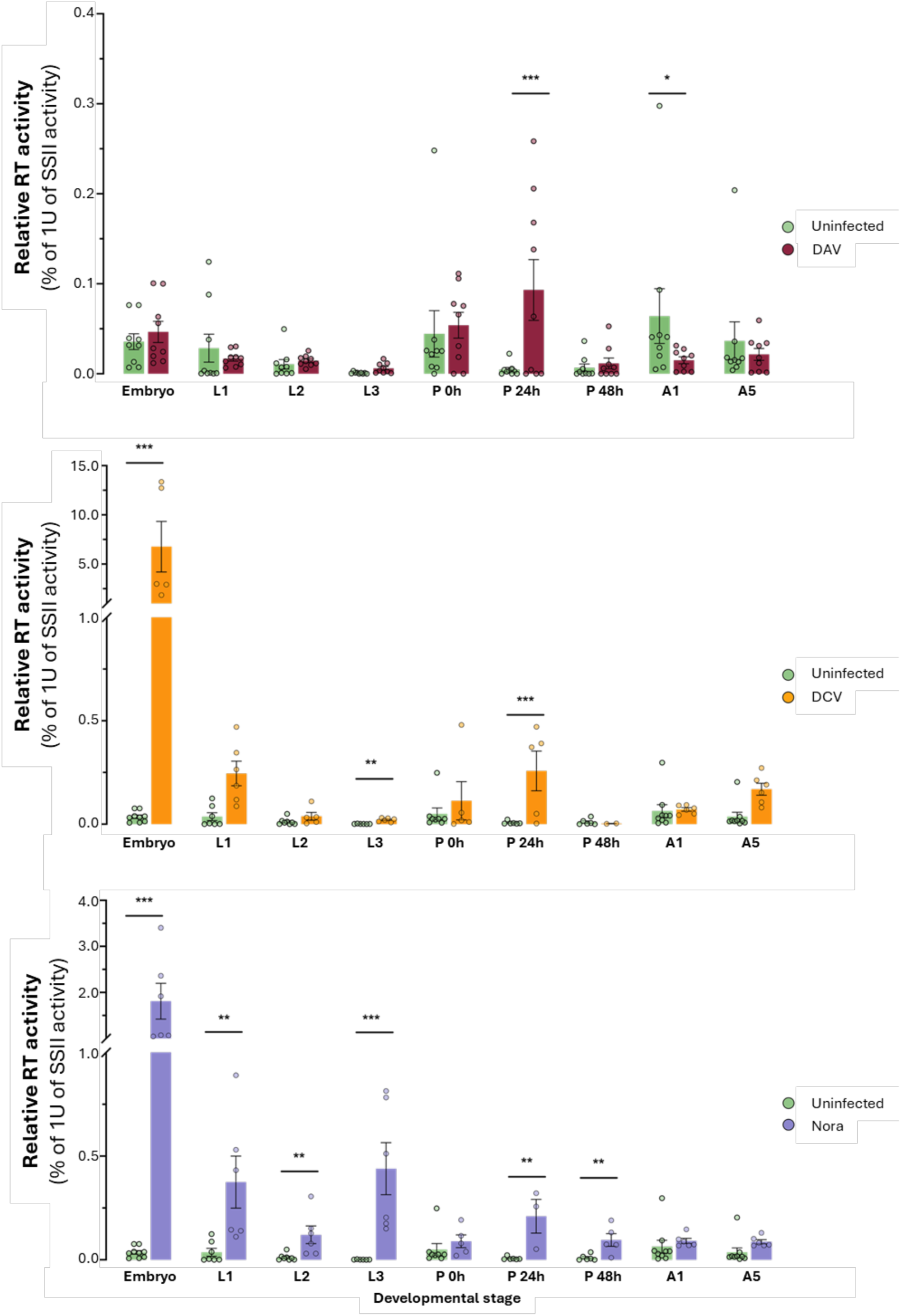
RT activity levels relative to 1 unit of SuperScript II across different developmental stages of infected or uninfected w^1118^ D. melanogaster stocks. In all panels, colors indicate infection status: green (uninfected), dark red (DAV), yellow (DCV), and light purple (Nora virus). A) From top to bottom, all infected stocks (DAV, DCV, and Nora virus) were compared to relative RT activity levels in the Uninfected stock. RT activities in infected versus Uninfected groups were compared within each developmental stage using generalized linear mixed models (GLMM, Gamma distribution) with condition-stage interaction and experiment random effects (P < 0.05, Tukey-adjusted pairwise comparisons). DAV, DCV, and Nora virus were analyzed separately.

**Supplementary Figure 6.**
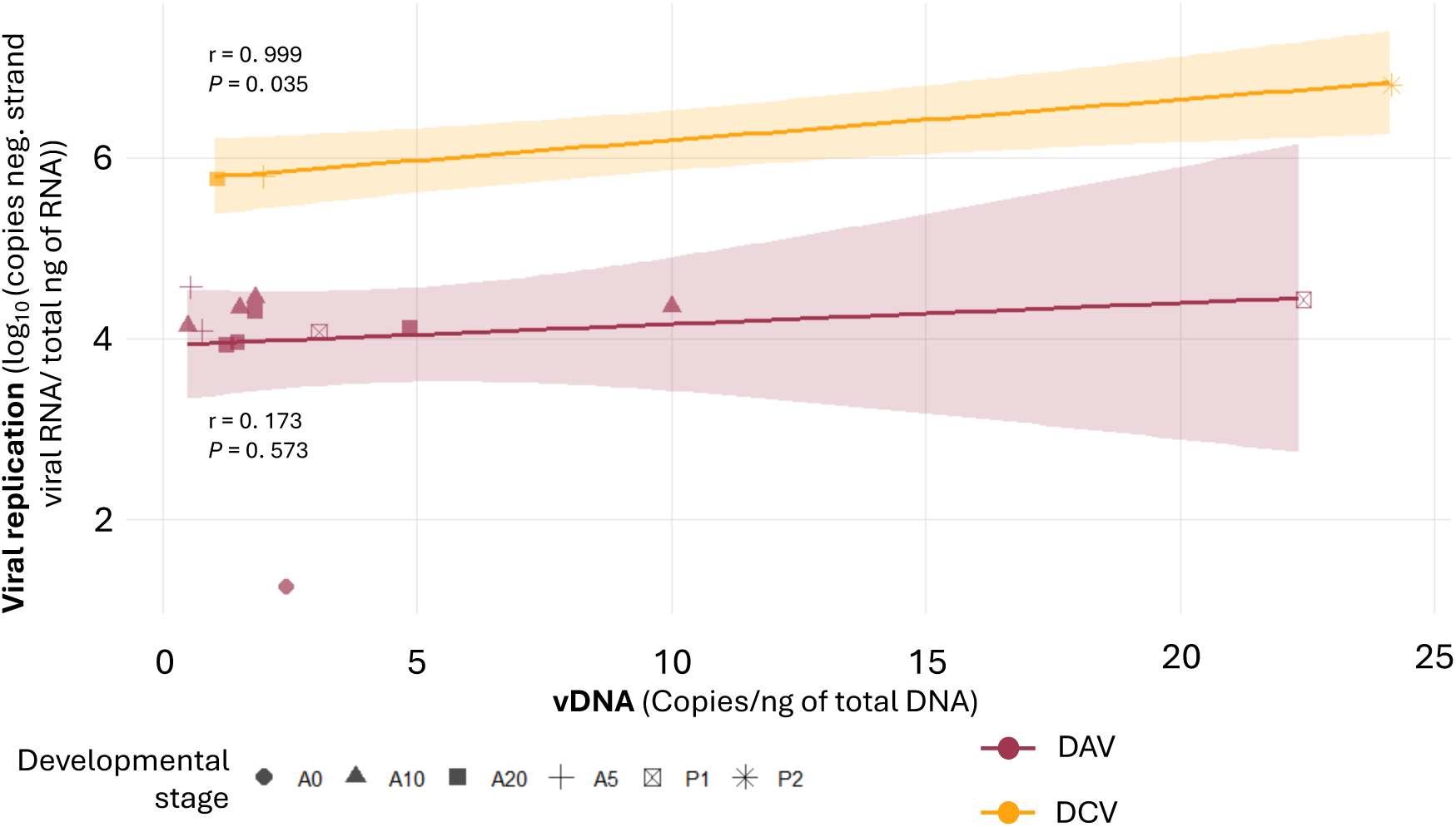
Viral replication levels and vDNA levels in DAV or DCV infected samples. Four samples were tested for vDNA for each stage and only samples in which vDNA was detectable by qPCR are shown. Y axis shows viral replication measured by RT-qPCR of the negative RNA strand, quantified using a standard curve. Viral replication is expressed as log10 of the copies of negative-strand viral RNA per total ng of RNA in each sample. X axis shows vDNA copies normalized to total DNA content (copies/ng). The R square value for each linear trendline is plotted. Dot shape represents the developmental stage, and dot color indicates the virus.

**Supplementary Figure 7.**
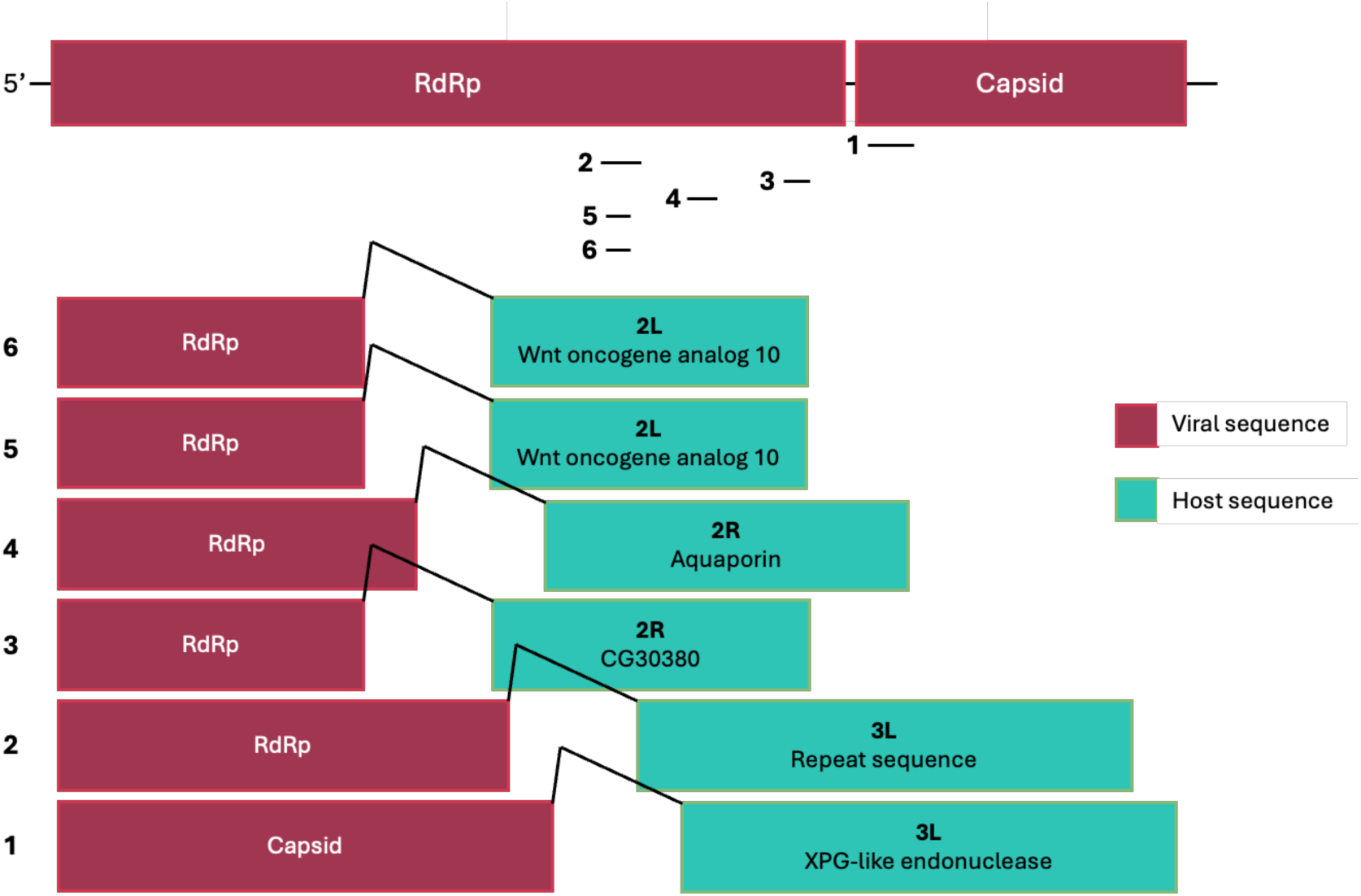
Chimeric virus-host reads from dsDNA sequencing. DNA extracted from DAV-infected pupae at 24 h post-pupation. The DAV genome is shown at the top of the schematic, followed by genomic positions of virus-host junctions (numbered 16). The compositions of the six chimeric reads representing viral gene segment (left) versus host chromosome/genetic element (right) are shown.

**Supplementary Figure 8.**
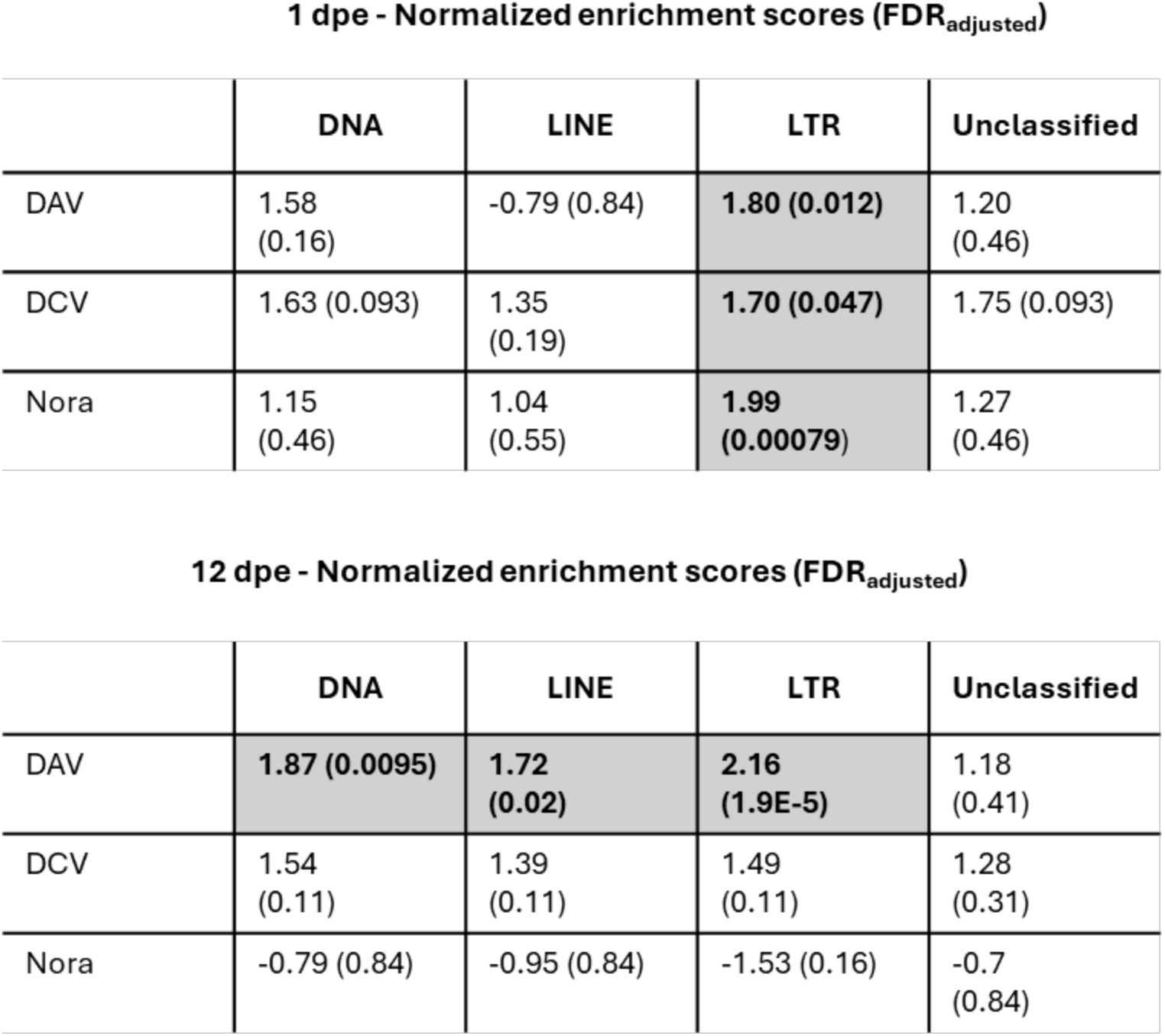
Normalized enrichment scores for each TE order in infected flies compared to uninfected flies. The adjusted False Discovery Rate (FDR) is shown between brackets. Cells highlighted in grey are those with a significant FDR. Top panel, 1 dpe adults. Bottom panel, 12 dpe adults.

---

**Supplementary Table 1.**
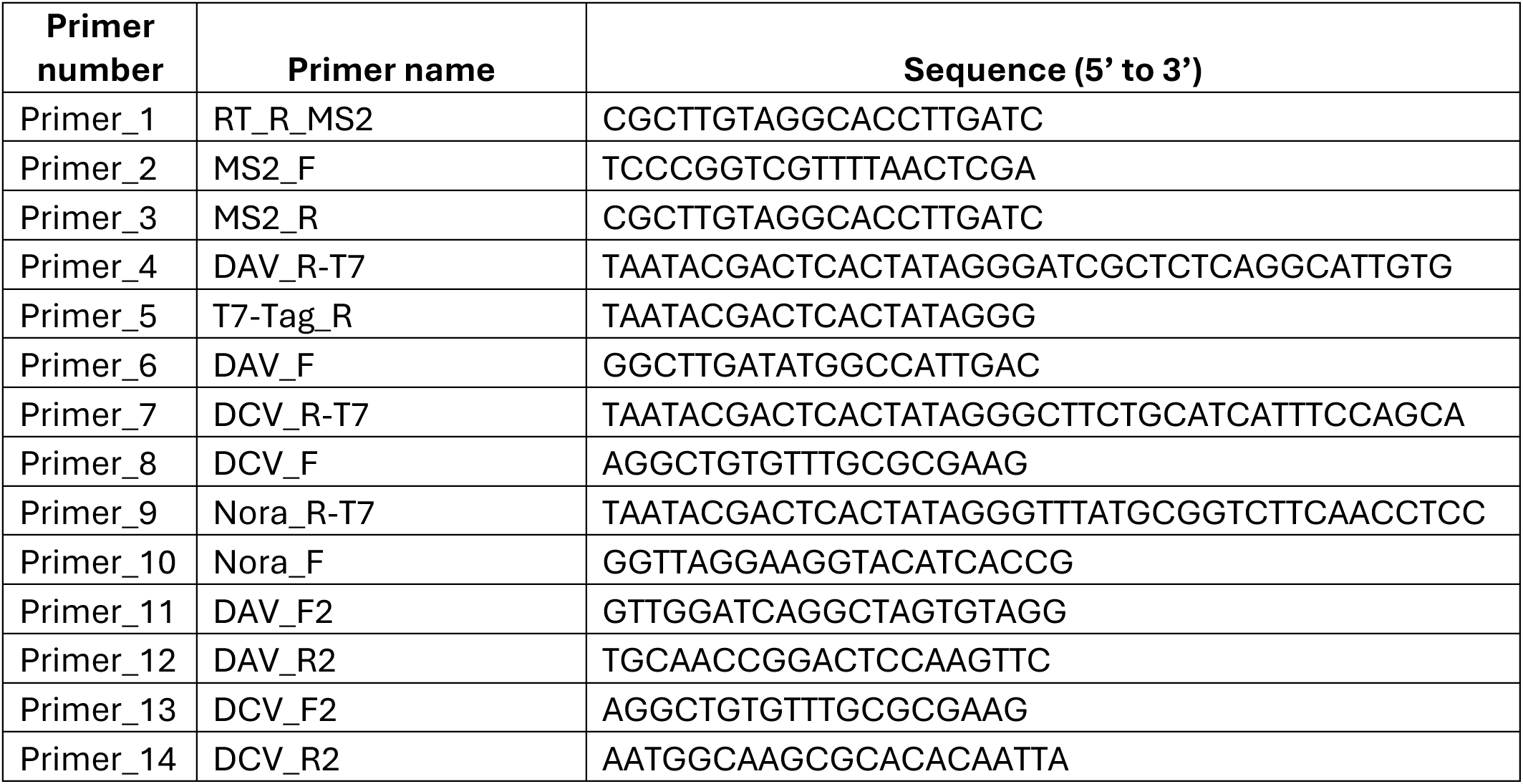
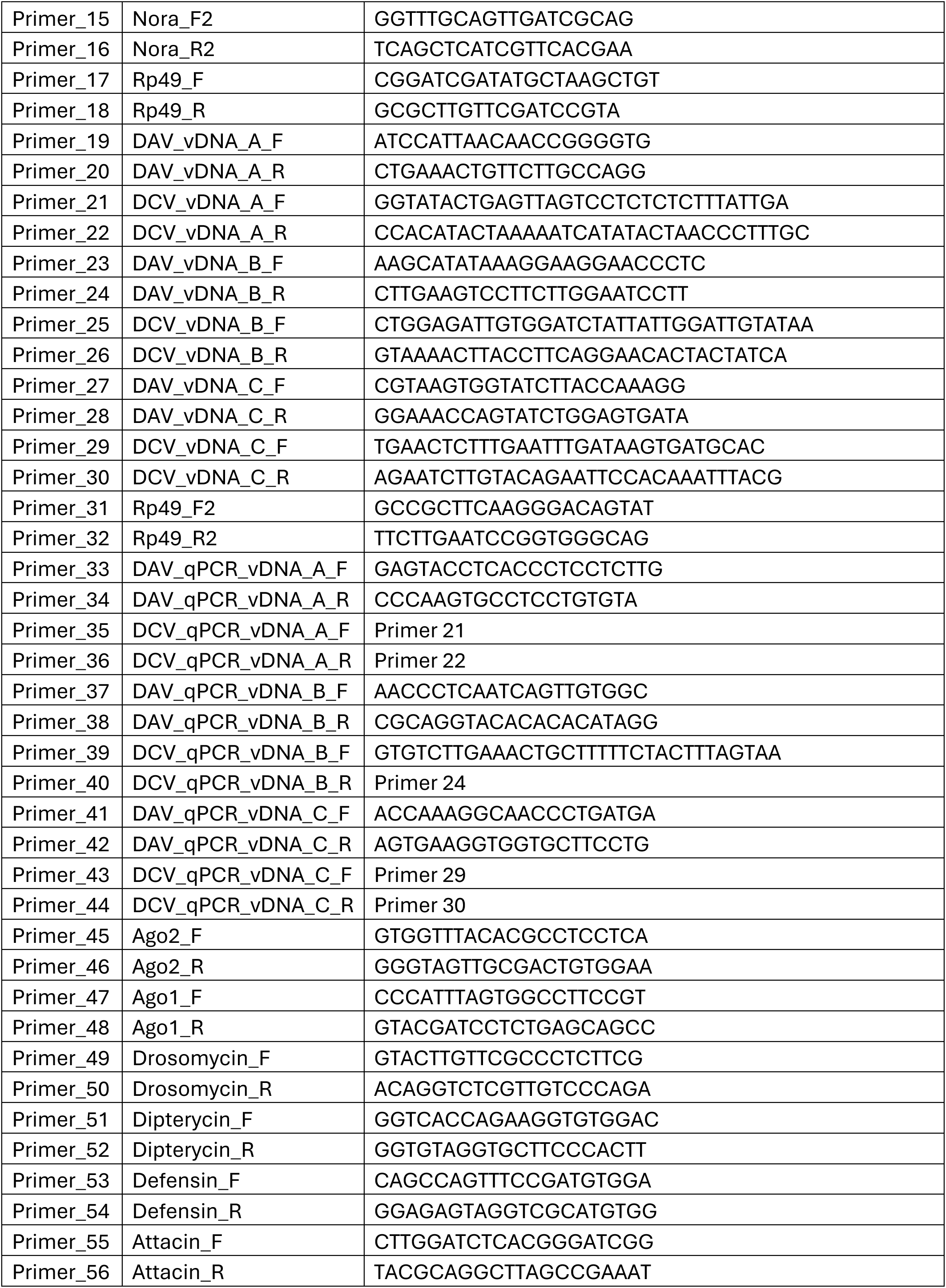
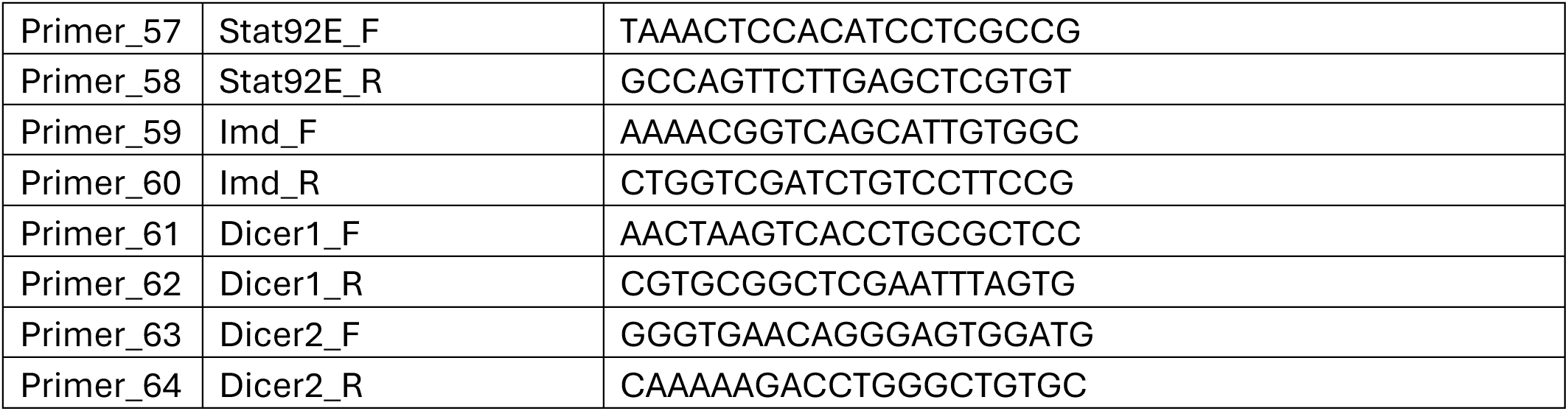
Primers used in this work. Each primer is identified with a number and a name, followed by the sequence.

**Supplementary Table 2.**
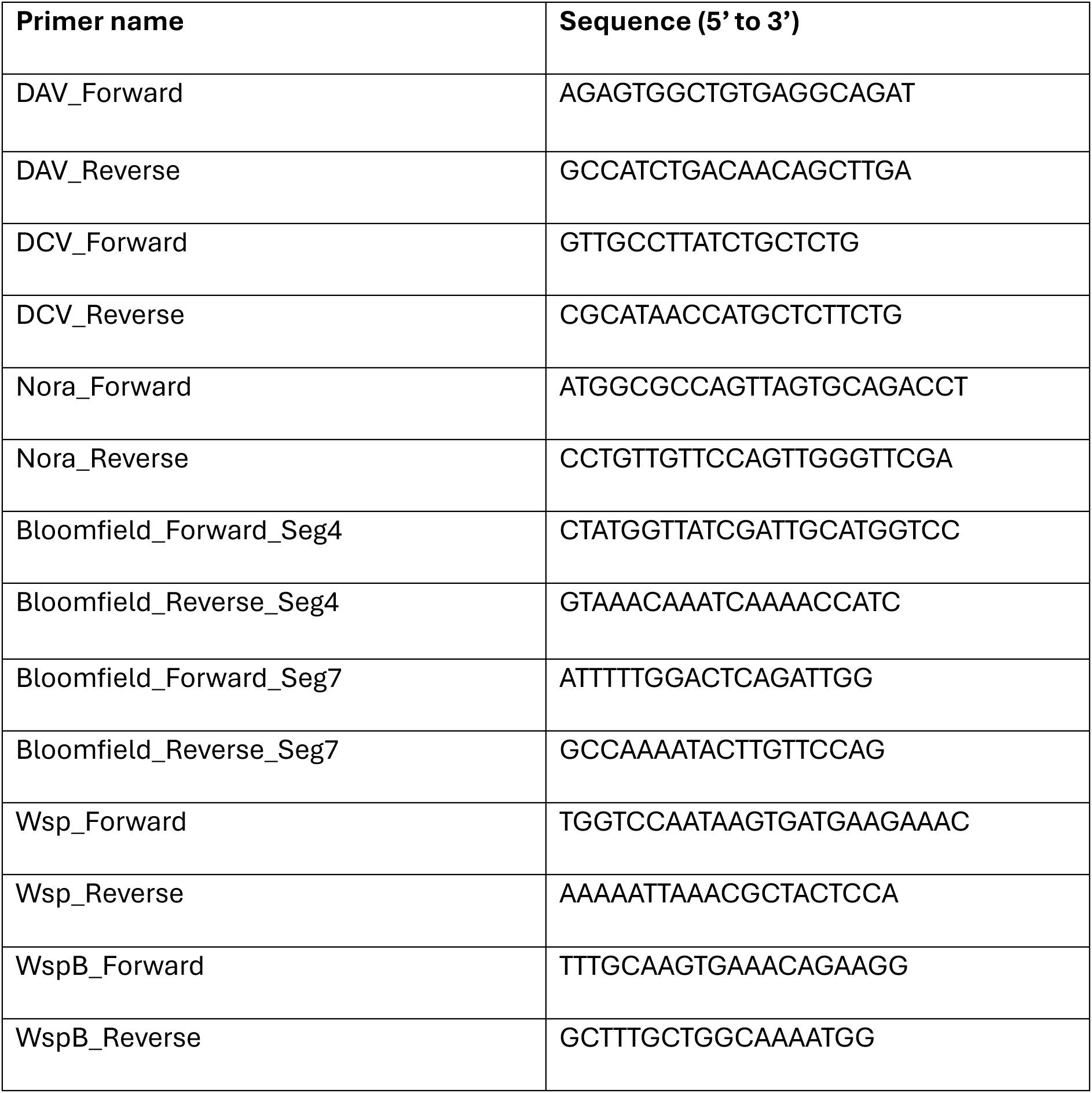
Primers used for RT-PCR to assess the presence of viruses and Wolbachia.

## References

Balakireva, Y., Nikitina, M., Makhnovskii, P., Kukushkina, I., Kuzmin, I., Kim, A., & Nefedova, L. (2024). The Lifespan of D. melanogaster Depends on the Function of the Gagr Gene, a Domesticated gag Gene of Drosophila LTR Retrotransposons. Insects, 15(1), 68. 10.3390/insects15010068

Banerjee, U., Girard, J. R., Goins, L. M., & Spratford, C. M. (2019). Drosophila as a Genetic Model for Hematopoiesis. Genetics, 211(2), 367–417. 10.1534/genetics.118.300223

Bergman, A., Crist, A. B., Lopez-Maestre, H., Blanc, H., Castelló-Sanjuán, M., Frangeul, L., Varet, H., Daron, J., Merkling, S. H., Saleh, M.-C., & Lambrechts, L. (2025). Limited impact of the siRNA pathway on transposable element expression in Aedes aegypti. BMC Biology, 23(1), 130. 10.1186/s12915-025-02225-8

Bonning, B. C., & Saleh, M.-C. (2021). The Interplay Between Viruses and RNAi Pathways in Insects. Annual Review of Entomology, 66(1), 61–79. 10.1146/annurev-ento-033020-090410

Castelló-Sanjuán, M., González, R., Romoli, O., Blanc, H., Nigg, J. C., & Saleh, M. C. (2025). Persistent viral infections impact key biological traits in Drosophila melanogaster. PLoS Biology, 23(10), 10.1371/journal.pbio.3003437.

Delahaye, C., & Nicolas, J. (2021). Sequencing DNA with nanopores: Troubles and biases. PLOS ONE, 16(10), e0257521. 10.1371/journal.pone.0257521

Dobin, A., Davis, C. A., Schlesinger, F., Drenkow, J., Zaleski, C., Jha, S., Batut, P., Chaisson, M., & Gingeras, T. R. (2013). STAR: Ultrafast universal RNA-seq aligner. Bioinformatics (Oxford, England), 29(1), 15–21. 10.1093/bioinformatics/bts635

Franklinos, L. H. V., Jones, K. E., Redding, D. W., & Abubakar, I. (2019). The effect of global change on mosquito-borne disease. The Lancet Infectious Diseases, 19(9), e302– e312. 10.1016/S1473-3099(19)30161-6

Gammon, D. B., & Mello, C. C. (2015). RNA interference-mediated antiviral defense in insects. Current Opinion in Insect Science, 8, 111–120. 10.1016/j.cois.2015.01.006

Garambois, C., Boulesteix, M., & Fablet, M. (2024). Effects of Arboviral Infections on Transposable Element Transcript Levels in Aedes aegypti. Genome Biology and Evolution, 16(5), evae092. 10.1093/gbe/evae092

Goic, B., & Saleh, M.-C. (2012). Living with the enemy: Viral persistent infections from a friendly viewpoint. Current Opinion in Microbiology, 15(4), 531–537. 10.1016/j.mib.2012.06.002

Goic, B., Stapleford, K. A., Frangeul, L., Doucet, A. J., Gausson, V., Blanc, H., Schemmel-Jofre, N., Cristofari, G., Lambrechts, L., Vignuzzi, M., & Saleh, M.-C. (2016). Virus-derived DNA drives mosquito vector tolerance to arboviral infection. Nature Communications, 7(1), 12410. 10.1038/ncomms12410

Goic, B., Vodovar, N., Mondotte, J. A., Monot, C., Frangeul, L., Blanc, H., Gausson, V., Vera-Otarola, J., Cristofari, G., & Saleh, M.-C. (2013). RNA-mediated interference and reverse transcription control the persistence of RNA viruses in the insect model Drosophila. Nature Immunology, 14(4), 396–403. 10.1038/ni.2542

Goodier, J. L. (2016). Restricting retrotransposons: A review. Mobile DNA, 7(1), 16. 10.1186/s13100-016-0070-z

Goodier, J. L., Pereira, G. C., Cheung, L. E., Rose, R. J., & Kazazian, H. H. (2015). The Broad-Spectrum Antiviral Protein ZAP Restricts Human Retrotransposition. PLOS Genetics, 11(5), e1005252. 10.1371/journal.pgen.1005252

Hadley Wickham. (2016). ggplot2: Elegant Graphics for Data Analysis. Springer-Verlag New York. https://ggplot2.tidyverse.org

Huang, Q., Gavor, E., Tulsian, N. K., Fan, J., Lin, Q., Mok, Y. K., Kini, R. M., & Sivaraman, J. (2023). Structural and functional characterization of Aedes aegypti pupal cuticle protein that controls dengue virus infection. Protein Science, 32(10), e4761. 10.1002/pro.4761

Hub / RNAflow GitLab. (2024, February 9). GitLab. https://gitlab.pasteur.fr/hub/rnaflow

Jin, Y., Tam, O. H., Paniagua, E., & Hammell, M. (2015). TEtranscripts: A package for including transposable elements in differential expression analysis of RNA-seq datasets. Bioinformatics, 31(22), 3593–3599. 10.1093/bioinformatics/btv422

Johnston, P. R., Paris, V., & Rolff, J. (2019). Immune gene regulation in the gut during metamorphosis in a holo-versus a hemimetabolous insect. Philosophical Transactions of the Royal Society B: Biological Sciences, 374(1783), 20190073. 10.1098/rstb.2019.0073

Kim, V. N., Han, J., & Siomi, M. C. (2009). Biogenesis of small RNAs in animals. Nature Reviews Molecular Cell Biology, 10(2), 126–139. 10.1038/nrm2632

Korotkevich, G., Sukhov, V., Budin, N., Shpak, B., Artyomov, M. N., & Sergushichev, A. (2016). Fast gene set enrichment analysis. Bioinformatics. 10.1101/060012

Lee, W.-S., Webster, J. A., Madzokere, E. T., Stephenson, E. B., & Herrero, L. J. (2019). Mosquito antiviral defense mechanisms: A delicate balance between innate immunity and persistent viral infection. Parasites & Vectors, 12(1), 165. 10.1186/s13071-019-3433-8

Li-Byarlay, H., Boncristiani, H., Howell, G., Herman, J., Clark, L., Strand, M. K., Tarpy, D., & Rueppell, O. (2020). Transcriptomic and Epigenomic Dynamics of Honey Bees in Response to Lethal Viral Infection. Frontiers in Genetics, 11, 566320. 10.3389/fgene.2020.566320

Love, M. I., Huber, W., & Anders, S. (2014). Moderated estimation of fold change and dispersion for RNA-seq data with DESeq2. Genome Biology, 15(12), 550. 10.1186/s13059-014-0550-8

Merkling, S. H., & Van Rij, R. P. (2015). Analysis of resistance and tolerance to virus infection in Drosophila. Nature Protocols, 10(7), 1084–1097. 10.1038/nprot.2015.071

Mikhailov, V. S., Zemskov, E. A., & Abramova, E. B. (1992). Protein synthesis in pupae of the silkworm Bombyx mori after infection with nuclear polyhedrosis virus: Resistance to viral infection acquired during pupal period. Journal of General Virology, 73(12), 3195–3202. 10.1099/0022-1317-73-12-3195

Mondotte, J. A., Gausson, V., Frangeul, L., Blanc, H., Lambrechts, L., & Saleh, M.-C. (2018). Immune priming and clearance of orally acquired RNA viruses in Drosophila. Nature Microbiology, 3(12), 1394–1403. 10.1038/s41564-018-0265-9

Mondotte, J. A., Gausson, V., Frangeul, L., Suzuki, Y., Vazeille, M., Mongelli, V., Blanc, H., Failloux, A.-B., & Saleh, M.-C. (2020). Evidence For Long-Lasting Transgenerational Antiviral Immunity in Insects. Cell Reports, 33(11), 108506. 10.1016/j.celrep.2020.108506

Nag, D. K., Brecher, M., & Kramer, L. D. (2016). DNA forms of arboviral RNA genomes are generated following infection in mosquito cell cultures. Virology, 498, 164–171. 10.1016/j.virol.2016.08.022

Nichols, C. D., Becnel, J., & Pandey, U. B. (2012). Methods to assay Drosophila behavior. Journal of Visualized Experiments: JoVE, 61, 3795. 10.3791/3795

Nigg, J. C., Castelló-Sanjuán, M., Blanc, H., Frangeul, L., Mongelli, V., Godron, X., Bardin, A. J., & Saleh, M.-C. (2024). Viral infection disrupts intestinal homeostasis via Sting-dependent NF-κB signaling in Drosophila. Current Biology, 34(13), 2785–2800.e7. 10.1016/j.cub.2024.05.009

Nigg, J. C., Mongelli, V., Blanc, H., & Saleh, M.-C. (2022). Innovative Toolbox for the Quantification of Drosophila C Virus, Drosophila A Virus, and Nora Virus. Journal of Molecular Biology, 434(6), 167308. 10.1016/j.jmb.2021.167308

Nunes, C., Koyama, T., & Sucena, É. (2021). Co-option of immune effectors by the hormonal signalling system triggering metamorphosis in Drosophila melanogaster. PLOS Genetics, 17(11), e1009916. 10.1371/journal.pgen.1009916

Öztürk-Çolak, A., Marygold, S. J., Antonazzo, G., Attrill, H., Goutte-Gattat, D., Jenkins, V. K., Matthews, B. B., Millburn, G., Dos Santos, G., Tabone, C. J., FlyBase Consortium, Perrimon, N., Gelbart, S. R., Broll, K., Crosby, M., Dos Santos, G., Falls, K., Gramates, L. S., Jenkins, V. K.,…Lovato, T. (2024). FlyBase: Updates to the Drosophila genes and genomes database. GENETICS, 227(1), iyad211. 10.1093/genetics/iyad211

Pino-Jiménez, B., Giannios, P., & Casanova, J. (2023). Polyploidy-associated autophagy promotes larval tracheal histolysis at Drosophila metamorphosis. Autophagy, 19(11), 2972–2981. 10.1080/15548627.2023.2231828

Poirier, E. Z., Goic, B., Tomé-Poderti, L., Frangeul, L., Boussier, J., Gausson, V., Blanc, H., Vallet, T., Loyd, H., Levi, L. I., Lanciano, S., Baron, C., Merkling, S. H., Lambrechts, L., Mirouze, M., Carpenter, S., Vignuzzi, M., & Saleh, M.-C. (2018). Dicer-2-Dependent Generation of Viral DNA from Defective Genomes of RNA Viruses Modulates Antiviral Immunity in Insects. Cell Host & Microbe, 23(3), 353–365.e8. 10.1016/j.chom.2018.02.001

Pyra, H., Böni, J., & Schüpbach, J. (1994). Ultrasensitive retrovirus detection by a reversetranscriptase assay based on product enhancement. Proceedings of the National Academy of Sciences, 91(4), 1544–1548. 10.1073/pnas.91.4.1544

Randall, R. E., & Griffin, D. E. (2017). Within host RNA virus persistence: Mechanisms and consequences. Current Opinion in Virology, 23, 35–42. 10.1016/j.coviro.2017.03.001

Rodriguez-Andres, J., Axford, J., Hoffmann, A., & Fazakerley, J. (2024). Mosquito transgenerational antiviral immunity is mediated by vertical transfer of virus DNA sequences and RNAi. iScience, 27(1), 108598. 10.1016/j.isci.2023.108598

Roy, M., Viginier, B., Saint-Michel, É., Arnaud, F., Ratinier, M., & Fablet, M. (2020). Viral infection impacts transposable element transcript amounts in Drosophila. Proceedings of the National Academy of Sciences, 117(22), 12249–12257. 10.1073/pnas.2006106117

Santos, D., Mingels, L., Vogel, E., Wang, L., Christiaens, O., Cappelle, K., Wynant, N., Gansemans, Y., Van Nieuwerburgh, F., Smagghe, G., Swevers, L., & Vanden Broeck, J. (2019). Generation of Virus-and dsRNA-Derived siRNAs with Species-Dependent Length in Insects. Viruses, 11(8), 738. 10.3390/v11080738

Salvati, M. V., Salaris, C., Monteil, V., Del Vecchio, C., Palù, G., Parolin, C., Calistri, A., Bell-Sakyi, L., Mirazimi, A., & Salata, C. (2021). Virus-Derived DNA Forms Mediate the Persistent Infection of Tick Cells by Hazara Virus and Crimean-Congo Hemorrhagic Fever Virus. Journal of Virology, 95(24), e01638–21. 10.1128/JVI.01638-21

Siomi, H., & Siomi, M. C. (2009). On the road to reading the RNA-interference code. Nature, 457(7228), 396–404. 10.1038/nature07754

Sivaprakash, M., Bhat, M. R., Bora, N. R., Patil, P. B., Sivakumar, B., & Rajah, R. A. (2025). Transmission Dynamics of Bombyx mori Nuclear Polyhedrosis Virus and Its Impact on the Economic Traits of Silkworm, Bombyx mori Linn. Journal of the Kansas Entomological Society, 98(1). 10.2317/0022-8567-98.1.10

Tassetto, M., Kunitomi, M., & Andino, R. (2017). Circulating Immune Cells Mediate a Systemic RNAi-Based Adaptive Antiviral Response in Drosophila. Cell, 169(2), 314-325.e13. 10.1016/j.cell.2017.03.033

Tettamanti, G., & Casartelli, M. (2019). Cell death during complete metamorphosis. Philosophical Transactions of the Royal Society B: Biological Sciences, 374(1783), 20190065. 10.1098/rstb.2019.0065

Treangen, T. J., & Salzberg, S. L. (2011). Repetitive DNA and next-generation sequencing: Computational challenges and solutions. Nature Reviews. Genetics, 13(1), 36–46. 10.1038/nrg3117

Trivedi, S., & Starz-Gaiano, M. (2018). Drosophila Jak/STAT Signaling: Regulation and Relevance in Human Cancer and Metastasis. International Journal of Molecular Sciences, 19(12), 4056. 10.3390/ijms19124056

Ullah, A., Tlak Gajger, I., Majoros, A., Dar, S. A., Khan, S., Kalimullah, Haleem Shah, A., Nasir Khabir, M., Hussain, R., Khan, H. U., Hameed, M., & Anjum, S. I. (2021). Viral impacts on honey bee populations: A review. Saudi Journal of Biological Sciences, 28(1), 523–530. 10.1016/j.sjbs.2020.10.037

Ullastres, A., Merenciano, M., & González, J. (2021). Regulatory regions in natural transposable element insertions drive interindividual differences in response to immune challenges in Drosophila. Genome Biology, 22(1), 265. 10.1186/s13059-021-02471-3

Wang, L., Tracy, L., Su, W., Yang, F., Feng, Y., Silverman, N., & Zhang, Z. Z. Z. (2022). Retrotransposon activation during Drosophila metamorphosis conditions adult antiviral responses. Nature Genetics, 54(12), 1933–1945. 10.1038/s41588-022-01214-9

Wichroski, M. J., Robb, G. B., & Rana, T. M. (2006). Human Retroviral Host Restriction Factors APOBEC3G and APOBEC3F Localize to mRNA Processing Bodies. PLoS Pathogens, 2(5), e41. 10.1371/journal.ppat.0020041

Wu, J., Wu, C., Xing, F., Cao, L., Zeng, W., Guo, L., Li, P., Zhong, Y., Jiang, H., Luo, M., Shi, G., Bu, L., Ji, Y., Hou, P., Peng, H., Huang, J., Li, C., & Guo, D. (2021). Endogenous reverse transcriptase and RNase H-mediated antiviral mechanism in embryonic stem cells. Cell Research, 31(9), 998–1010. 10.1038/s41422-021-00524-7

Zhang, Y., Zhang, X., Dai, K., Zhu, M., Liang, Z., Pan, J., Zhang, Z., Xue, R., Cao, G., Hu, X., & Gong, C. (2022). Bombyx mori Akirin hijacks a viral peptide vSP27 encoded by BmCPV circRNA and activates the ROS-NF-κB pathway against viral infection. International Journal of Biological Macromolecules, 194, 223–232. 10.1016/j.ijbiomac.2021.11.201

Zattera, M. L., & Bruschi, D. P. (2022). Transposable Elements as a Source of Novel Repetitive DNA in the Eukaryote Genome. Cells, 11(21), 3373. 10.3390/cells11213373

Zeng, M., Yan, Z.-Y., Lv, Y.-N., Zeng, J.-M., Ban, N., Yuan, D.-W., Li, S., Luan, Y.-X., & Bai, Y. (2025). Molecular basis of E93-dependent tissue morphogenesis and histolysis during insect metamorphosis. Insect Biochemistry and Molecular Biology, 177, 104249. 10.1016/j.ibmb.2024.104249

